# Rare non-synonymous germline mutations systematically define the risk of triple negative breast cancer

**DOI:** 10.1101/302398

**Authors:** Mei Yang, Yanhui Fan, Zhi-Yong Wu, Zhendong Feng, Qiangzu Zhang, Shunhua Han, Xiaoling Li, Teng Zhu, Minyi Cheng, Juntao Xu, Ciqiu Yang, Hongfei Gao, Chunming Zhang, Guangming Tan, Michael Q. Zhang, You-Qiang Song, Gang Niu, Kun Wang

**Author notes:** These authors contributed equally to the work (Mei Yang, Yanhui Fan, Zhi-Yong Wu). Corresponding Author: Gang Niu, Phil Rivers Technology, Beijing, 100095, China.; and Kun Wang, Guangdong Provincial People’s Hospital, Guangdong Academy of Medical Sciences, Guangzhou, 510080, China.

## Abstract

Early identification of the risk for triple-negative breast cancer (TNBC) at the asymptomatic phase could lead to better prognosis. Here we developed a machine learning method to quantify systematic impact of all rare germline mutations on each pathway. We collected 106 TNBC patients and 287 elder healthy women controls. The spectra of activity profiles in multiple pathways were mapped and most pathway activities exhibited globally suppressed by the portfolio of individual germline mutations in TNBC patients. Accordingly, all individuals were delineated into two types: A and B. Type A patients could be differentiated from controls (AUC = 0.89) and sensitive to BRCA1/2 damages; Type B patients can be also differentiated from controls (AUC = 0.69) but probably being protected from BRCA1/2 damages. Further we found that Individuals with the lowest activity of selected pathways had extreme high relative risk (up to 21.67 in type A) and increased lymph node metastasis in these patients. Our study showed that genomic DNA contains information of unimaginable pathogenic factors. And this information is in a distributed form that could be applied to risk assessment for more cancer types.

**Significance:** We identified individuals who are more susceptible to triple negative breast cancer. Our method performs much better than previous assessments based on BRCA1/2 damages, even polygenic risk scores. We disclosed previously unimaginable pathogens in a distributed form on genome and extended risk prediction to scenarios for other cancers.

## Introduction

The continuously increasing prevalence and mortality of breast cancers is a major health concern in women worldwide (1–3). Triple-negative breast cancer (TNBC) is a breast cancer subtype with absence of estrogen receptor (ER), progesterone receptor (PR) and human epidermal growth factor 2 (HER2), which makes up 15% to 20% of all breast cancer cases (4). TNBC is clinically aggressive because 1) the average age of initial diagnosis is 4 - 13 years earlier compared to other types of breast cancer (5–12); 2) The five-year survival rate is low, which reflects fast progression and poor prognosis (13–15); 3) lack of efficient therapy strategies due to the lack of either endocrine or anti-HER2 targets (12,16,17). Nonetheless, identification of the biomarkers for prognosis at earlier stages including the asymptomatic stage could facilitate timely treatment and expand the surgery window for breast cancer patients - TNBC the most, as in reducing the mortality (18,19).

Evidence that supports genetic hereditary features of breast cancer, especially TNBC, forms the foundation of our rationale. Inherited mutations have been observed in multiple epidemiological studies and clinical trials of TNBC with a positive percentage generally higher than 35% across different ethnicities (5,9,12,20,21), where the hereditary predisposition for sporadic cases might be masked by a limited family structures and driven in a non-Mendelian fashion (22). High penetrance predisposition genes, such as the notorious BRCA1/2 and other genes that function in homologous recombination (HR) repairing of double strand DNA breaks (23–28), provide more profound evidence for the germline heredity of TNBC. BRCA1/2 germline mutations are found in 5.3% of unselected breast cancers according to The Cancer Genome Atlas (TCGA) while the prevalence percentages rise up to 9.4 to 18.2% in TNBC from different study cohorts (5,7–10,29–32). In addition, data from investigations of genetic counselling showed that 40% of BRCA1/2 germline mutation carriers have been diagnosed with TNBC before the age of 40, twice as high as the proportion in all unselected breast cancer cases (12,33). These findings suggest that the germline mutations might play a more important role in TNBC compared to other types of breast cancer, which might contribute to its early onset, as well as other features.

However, germline determined risk is far from being explicitly evaluated in TNBC breast cancer. On the aspect of population coverage, the prevalence of disease-associated mutations for each individual gene is generally less than 5% except for BRCA1/2 (23,25,27,28), making it hard to extract the distribution of common features in heterogeneous disease such as TNBC with merely one or several genes. On the other hand, current predisposition genes are mostly limited in homologous recombination genes identified from studies based on different cohorts that report diverged or even controversial results especially when interventions, such as therapies, are introduced (5,34–39). These all suggest variable features of connections within the disease are not described by the known genes and thus call for more holistic evaluation approaches.

Quantified genome-wide polygenic approaches have been adopted in predicting risks of multiple genetic diseases, such as the coronary artery disease and Type 2 diabetes (40), and the deep learning algorithms are adopted to empower robust data-driven feature recognition in the application of identifying human pathogenic germline mutations (41) as well as gene expression projection from the genome mutations (42). Nonetheless, these models’ performance egregiously depends on the degree of inherent heterogeneity of diseases, meaning the fewer numbers of biological functions are affected by the disease, the better their representations would be (40–42). Since cancers are significantly more heterogeneous than other complex genetic diseases, the applications of previous models have been constrained (14). Our study aims to tackle a question that necessitates an answer yet not solved - who are more likely to suffer from the triple-negative breast cancer? Our goal is to find a quantitative standard using the germline mutation information - the earliest predictive features in any given individual, so that susceptible individual could be accurately identified from the female population with false negatives and false positives reduced at the same time. The specific approach we take is primarily to build a machine learning framework based on genetic data and prior information about biological functions. The phylogenetic coding variation information is projected step by step in a new spatial space that reflects the activity of pathways. Based on this, we find that healthy risk-free individuals are significantly spatially separated from patients and all the subjects can be clustered according to their distributions. Intriguingly, we find the identified classifications could predict relative risk of developing TNBC, clinical staging characteristics; and the pathogenicity of BRCA1/2 is associated with the identified classifications, supporting the validity of the methodology and its future application to a wide range of disease risk assessment scenarios.

## Materials and Methods

### Study participants

The study cohorts included 106 TNBC patients recruited from this study and 287 cancer-free elderly women (CTRL) over 80 years old from an open dataset (43), which served as the control. The statistics of clinical and general information were shown in supplementary Table 1-3 for TNBC patients and supplementary Table 4 for the Control. The study protocol was approved by the Research Ethics Committee at Guangdong General Hospital, Guangdong Academy of Medicals Sciences. Written informed consent was obtained from all participants for the use of banked tissues (including white blood cells and buccal cells), collection of pathological data and clinical follow-up data. The SNP data on CTRL was gathered from a previous report (43).

### Whole-exome sequencing and mutation calling

Peripheral blood mononuclear cells (PBMC) were collected from TNBC patients, which was defined by ≤1% estrogen receptor (ER) positive tumor cells, progesterone receptor (PR) negativity, and normal HER2-receptor expression (44). Each sample was prepared according to the Illumina protocols. Paired-end multiplex sequencing of samples was performed on an Illumina HiSeq X Ten sequencing platform. Samples from the CTRL cohort were sequenced on an Illumina HiSeq 2000 system. TNBC cases were sequenced to a median depth > 120X and controls were sequenced to a median depth > 60X. Paired-end raw sequence reads were mapped to the human reference genome (UCSC hg19) using the Burrows-Wheeler Aligner (45) with default settings. Mutation calling was carried out using the HaplotypeCaller module in the Genome Analysis Toolkit (GATK) (46) according to GATK Best Practices. Briefly, the aligned BAM files were first marked for duplicated reads by Picard. Local realignment around indels and base quality score recalibration were performed by GATK. The processed BAM files were then used to call SNPs and indels. Mutation filtering of SNPs and indels were performed separately by mutation quality score recalibration using GATK. The filtered mutations were then annotated by ANNOVAR (47).

### Statistical analysis

The R package *pheatmap* was used to plot the clustered heatmaps using Ward’s criteria (48). Naïve Bayes implemented in R package caret was performed to build the binary classifiers to separate the TNBC patients from the CTRL. All data were randomly split into the training group (60%) and the testing group (40%) preserving the overall class distribution of the data. The optimal parameters were selected by 5-fold cross validation. Several criteria including area under the receiver operating characteristics curve (AUC), accuracy, sensitivity, and specificity were used to measure the performance of the classifier. Accuracy was calculated as the proportion of true positives and true negatives in all evaluated cases, sensitivity was the probability of testing positive with the disease present, and specificity was the probability of testing negative with no disease present. The R package *ggscatter* was used to draw the scatter plots and linear regression lines with 95% confidence interval. Pearson correlation coefficients and p-values were also labeled on the plot. The NMRCLUST (49) method was used to cluster the pooled samples into an optimal number of classes. This method uses average linkage clustering followed by penalty functions to simultaneously optimize the number of clusters and the average spread of clusters. All statistical analyses and plots were conducted using in-house scripts developed in Perl, Python, R, and MATLAB.

### Data availability

The data that support this study are provided in supplementary tables. Whole exome sequencing data have been deposited in the Short Read Archive (SRA) under the BioProject ID PRJNA497746 (https://www.ncbi.nlm.nih.gov/sra/PRJNA497746). We have implemented our methods as a web server, which is freely available at http://philrivers.vicp.io:9900/.

## Results

### A machine learning model to delineate genome wide TNBC susceptibility

As populations tend to go through purifying selection for complex disease susceptibility mutations, we hypothesize that rare non-synonymous germline mutations are functional in TNBC susceptibility. The functional importance of these rare mutations, on the other hand, is hard to be delineated due to allelic heterogeneity and the possibility that they might work as a network collectively. We aspired to build a machine learning framework (Damage Assessment of Germline shotGun, DAGG) that projects the rare non-synonymous germline mutations of an individual in high-dimensional but sparse onto an informative low-dimensional spectrum of sixty signaling pathways, which have been concurrently reported in association with cancer, drug target or immune related terms in public knowledge databases.

DAGG included three components that work sequentially as illustrated in Fig. 1. The first component took the non-synonymous mutations and expression as input and the effect of each mutated gene on the expression of all genes as output. Genes were coded as 1 or 0 represent any mutations presented or not, respectively. Then a Z-score, defined as the driving force, to measure the statistical significance of causal connection from the mutations on DNA to RNA expression profiles was calculated for each gene using both the mutation and expression data from the COSMIC Cell Line Project (50). Briefly, for any given mutated gene *v*_*i*_, all cell lines were separated into two groups: *MT* group and *WT* group based on whether they carried the non-synonymous mutations in this gene *v*_*i*_. To investigate how the expression of any gene *g*_*j*_could be impacted under the given mutation profile of gene *v*_*i*_, the difference of the mean expression of gene *g*_*j*_ between the *MT* and the *WT* group was quantified and defined as the differential expression score Δ_*ij*_. To account for the background effect, we randomly group cell line while maintaining groups size for 100,000 times, the Z-score (as the driving force *F*_*ij*_) could be calculated with the following equation,

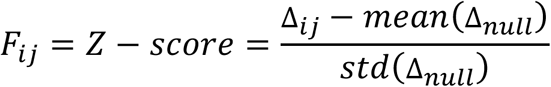

where Δ_*null*_ stands for the differential expression score for each random grouping while *mean* and *std* calculate their mean and standard deviation respectively. A bigger absolute value of *F*_*ij*_ represents stronger effect (called deterministic expression when |F_ij_| ≥ 3) of the mutated gene *v*_*i*_ on the expression of gene *g*_*j*_, and we took values either no smaller than 3 or no larger than −3 as the cutoff for significance. Traversing all the genes (assuming the total number of mutated genes is *m* and the total number of genes with expression data is *n*) through these steps would generate an *m x n* matrix of driving force (Fig.1B) and project the sparse mutation profile in a discrete space into a continuous space related to the gene expression profile, thus introducing the global gene regulation functions of non-synonymous mutations based on COSMIC cell line dataset in the first component.

**Fig. 1.**
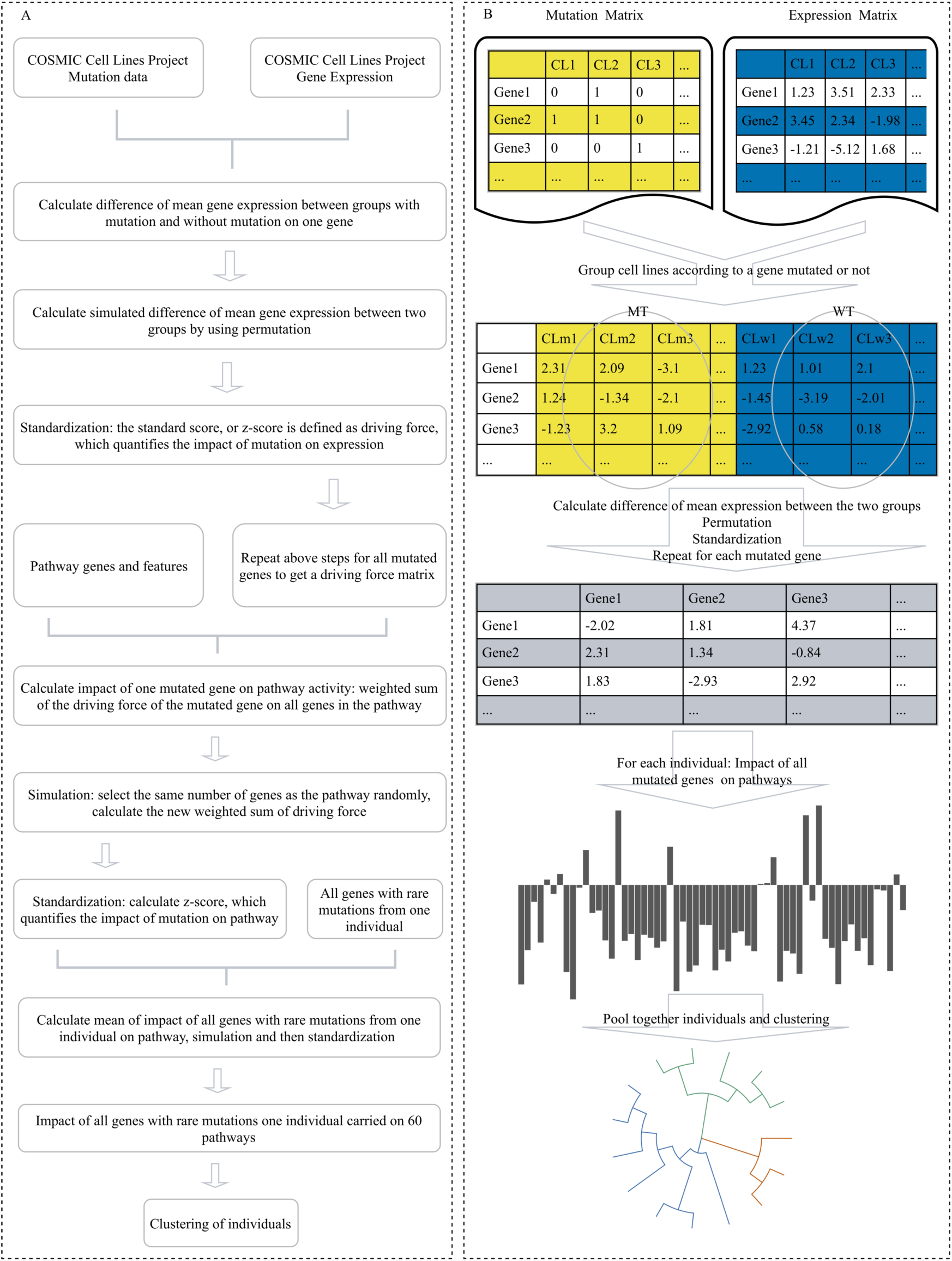
Workflow of our machine learning framework for constructing rare germline mutation-based pathway activity spectrum for TNBC/CTRL identification. (A) Workflow of the machine learning framework developed for this study. (B) Schematic overview of DAGG framework. Mutation and expression matrices are built for genes mutated in more than five cell lines using data from COSMIC Cell Lines Project (CCLP). For each mutated gene, the expression matrix was divided into two groups: mutant group (MT) and wildtype group (WT). The differentially expression score for each gene was calculated as the difference of mean expression of this gene between MT and WT groups. Then permutation was used to divided the expression matrix into two groups randomly while keep the number of cell lines in each group. We obtained the driving force for each mutated gene by calculating the z-score. Then impact of mutated genes on pathways were calculated. Finally, pooled samples of TNBC and CTRL were clustered into different classes and groups.

The second component translated the global changes in gene expression into activation or repression of signaling pathways. First, the effect of one mutated gene *v*_*i*_ on one pathway *P*_*j*_ activity was defined as the following equation:

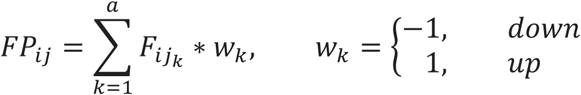

Where *F* is the driving force from the first component, *j*_*k*_ is the column position in the *m x n* matrix of the *k*_*th*_ gene in the pathway, *a* is the total number of genes involved in the *j*_*th*_ pathway, *w*_*k*_ is the weight of the *k*_*th*_ gene in the pathway. The weights for up- and down-regulated pathway signature were 1 and −1, respectively. The Z-score of *FP*_*ij*_ was calculated based on 100,000 times simulation in which a random gene set having the same number of genes as the specified pathway does was randomly selected for the calculation of the simulated *FP*. For a specific individual carried *b* mutated genes, the collective effect of these mutated genes on the pathway activity was defined as the summation of Z-score of *FP* of each mutated gene. Then another Z-score of the sum was calculated based on 100,000 times simulation in which *b* genes were randomly selected for the calculation of the simulated summation. The second component further reduces the high-dimensional sparse mutation data to features of sixty pathways (*1 x 60* matrixes compared to *m x n* in the first component) and made it easier to make comparisons among different individuals.

We hypothesized that the phenotypically similar individuals should have common pathway activity spectrum features which were adequately distinguishing between phenotypically differed cohorts, i.e. the TNBC and the healthy control. Therefore, we pre-selected pathways that were the most significantly impacted by rare non-synonymous germline mutations with Z-scores no larger than −4, and constructed the third component to assign each individual in pooled cohorts to specific classes and groups based on activity spectra of these pathways. Note the number of clusters determined by minimizing the spread across each cluster. The relative risk in each class was then calculated, followed by stratification of these classes into three groups by the range of relative risks and z-scores, where Group 1 was with the lowest combined relative risk and Group 3 the highest.

### TNBC patients significantly differentiated from healthy subjects by pathway activity spectrum

To lay the foundation, we first analyzed whole exome sequencing (WES) data of rare non-synonymous germline mutations from a cohort of 287 cancer free elderly women over 80 years old (287 subjects from LOAD cohort (43)) that used as the CTRL cohort, whom we postulate the germline genetics begets the natural resistant against TNBC to throughout evolution. After obtaining the features of their pathway spectra, we observed an intriguing phenomenon that all individuals exhibit either one of two typical differentiated pathway spectrum patterns that we identified as Type A and Type B (Fig. 2A). Among Type A subjects, the functional effect of the rare non-synonymous mutations could be clearly defined as up-regulation or down-regulation of the pathways. This means the functional effect of the mutations on the pathway activity was sufficiently significant to yield z-scores of large absolute values, so that clear cut-off could be set to define up- and down-regulation of pathways and generate high-contrast pattern on the heatmap. On the contrary, the functional effect of the rare mutations in Type B was ambiguous, demonstrating an overall reluctance against pathway activity alterations marked with low contrast patterns on the heatmap. These findings suggested an inherent difference in the “tolerance” of rare non-synonymous germline mutations among cancer free populations. Next for the TNBC cohort, we carried out WES of peripheral blood cells collected from 106 TNBC patients with a median depth > 120X in contrast to > 60X in the CTRL. Similarly, the TNBC patients also exhibited either Type A or Type B of overall pathway spectrum patterns (Fig. 2B). This finding indicates that the pathway spectrum typing is ubiquitous across populations as an inherent tolerance against rare non-synonymous germline mutations, potentially independent of the presence of TNBC. This also prompts that comparison of patients and healthy individuals should only be made after typing in order to rule out restrictions invited by the difference of tolerance against rare non-synonymous germline mutations. When we analyzed the pathway functional features within the Type A category, significant differences between CTRL and TNBC patients were identified from the heatmap in Fig. 2C, with TNBC patients well separated from the CTRL (AUC = 0.89) (Fig. 2D). Meanwhile, TNBC patients could also be separated from the CTRL in the Type B category (AUC = 0.69, Fig. 2E-F). These results suggest that global germline genetics plays a role in the susceptibility of TNBC, even though it is traditionally believed that the hereditary risk is relatively low for this type of cancer when germline mutations are reviewed individually.

**Fig. 2.**
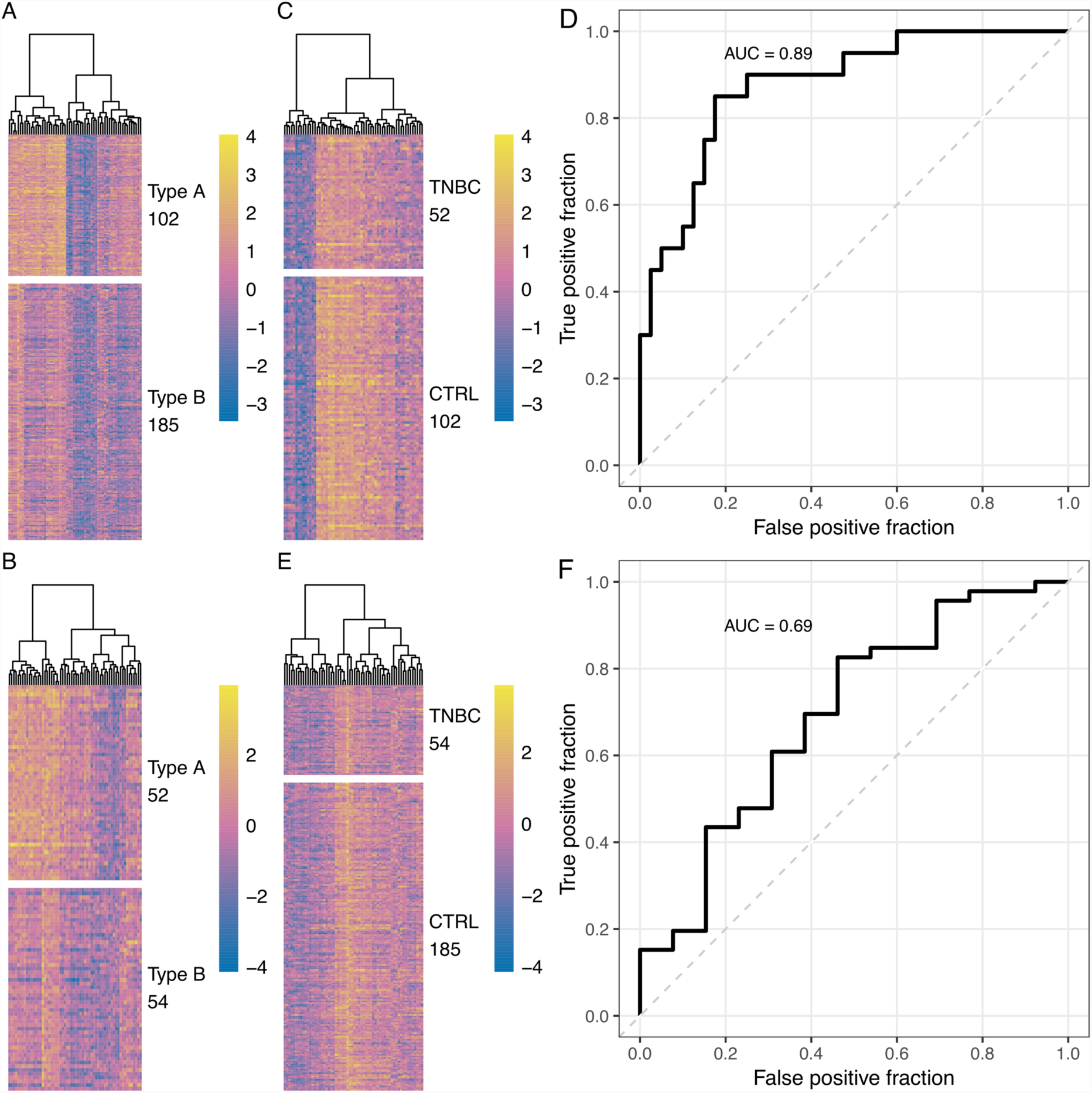
Individuals can be grouped into Type A and Type B. (A) Clustering heatmap of CTRL based on calculated pathway activity spectrums. Each row represents a CTRL individual and columns indicate 60 pathways. The population was characterized into two distinctive types, Type A and Type B. (B) Clustering heatmap of TNBC based on calculated pathway activity spectrums. Each row represents a TNBC individual and columns indicate 60 pathways. (C) Clustering heatmap of Type A TNBC and CTRL based on calculated pathway activity spectrums. Each row represents a TNBC or CTRL individual and columns indicate 60 pathways. (D) The receiver operating characteristic (ROC) curve for the classification performance in Type A. (E) Clustering heatmap of Type B TNBC and CTRL based on calculated pathway activity spectrums. Each row represents a TNBC or CTRL individual and columns indicate 60 pathways. (F) The receiver operating characteristic (ROC) curve for the classification performance in Type B.

### Functional pathway features coded by TNBC germline rare non-synonymous mutations

To mine the functional features specific to TNBC, we used the 52 TNBC patients and 102 CTRL in type A to compare the sixty pathways activities between the two cohorts. We calculated the differences in each pathway by subtracting the averaged values of a pathway in CTRL from that in TNBC. Next, we conducted 100 million times of permutation. Each permutation randomly shuffles the label of CTRL and TNBC for each subject so that the sample size of each group remains unchanged. Then the significance was calculated as the standard score that measures how many standard deviations deviated from the population mean (defined by Z-score). The results revealed that only 11 out of the 60 pathways had been positively regulated (3 of them are significantly regulated, Z-score ≥ 3), whereas the other 49 had been negatively regulated (25 pathways are significantly regulated, Z-score ≤ −3) (Fig. 3A-B). With approximately 87% signaling pathways suppressed in the activity, these results provide evidence that the germline genetics in TNBC patients tends to increase the vulnerability via suppression over the global pathway activity instead of over-activation on selected pathways.

**Fig. 3.**
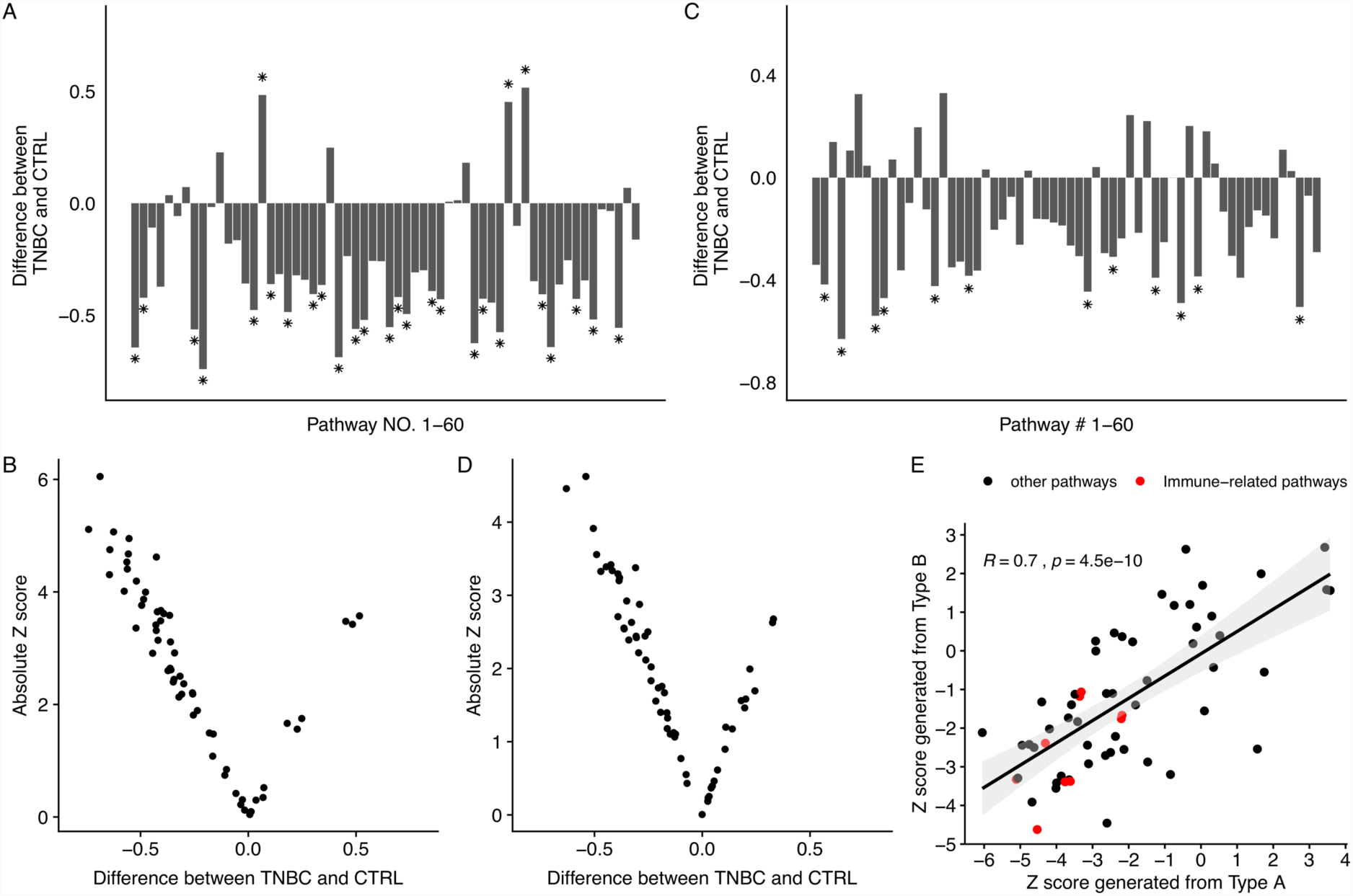
Downregulated pathways characterized Type A and Type B TNBC patients. (A) States of pathways in Type A. Differences larger than 0 represent upregulated pathways, whereas differences less than 0 represent downregulated pathways. Stars denote absolute values of Z scores of the differences larger than 3. (B) Scatterplot of absolute values of Z scores for the 60 pathways in Type A. (C) States of pathways in Type B. (D) Scatterplot of absolute values of Z scores for the 60 pathways in Type B. (E) The correlation of Z scores between Type A and Type B. The linear regression curve is shown as a black line and the shadowed area indicates the 95% CI.

Similar analyses were carried out for Type B CRTL and TNBC (Fig. 3C-D), and the global suppression on pathways (43 out of 60) was still a major event in terms of the down-regulation, despite only 12 pathways had significance with z-score ≤ −3. Although 17 pathways were defined as up-regulated in Type B, as opposed to 11 pathways in Type A, none of these pathways had sufficient significance with Z-score ≥ 3. Nonetheless, the pathway activity difference between TNBC patients and correspondent CTRL demonstrated similarities between Type A and Type B, giving a fitness of linear regression with r = 0.7 and p = 4.5e-10 for their functional alterations (Fig. 3E). This suggests that the functional pathway deviation along with TNBC development was conserved no matter what the starting pattern typing is, i.e. Type A or Type B. In addition, immune-related pathways were commonly down-regulated in both Type A and Type B (Z scores are all smaller than 0, Red dot in Fig. 3E).

### Activities of selected pathways define relative risk of TNBC

We hypothesize that a more detailed classification will be helpful to better determine the TNBC susceptibility from general population. Therefore, we applied a penalty model (51), which provided us an optimal total number of classes in each type, to the combined population of TNBC patients and CTRL subjects. The optimal number of classes was fifteen for Type A and thirteen for Type B (Fig. 4A-B). The classes of Type A and those of Type B were presented by eight and six selected pathways (Z-score <= −4), respectively. TNBC patients and CTRL subjects were not uniformly distributed among different classes (Table 1 and Table 2). The odds of TNBC patients to CTRL individuals was calculated to evaluate the relative risks of each class, where classes with more TNBC patients were esteemed as of high relative risk and classes with more CTRL subjects as of low relative risk plus classes with approximately even proportions as of moderate relative risk.

**Fig. 4.**
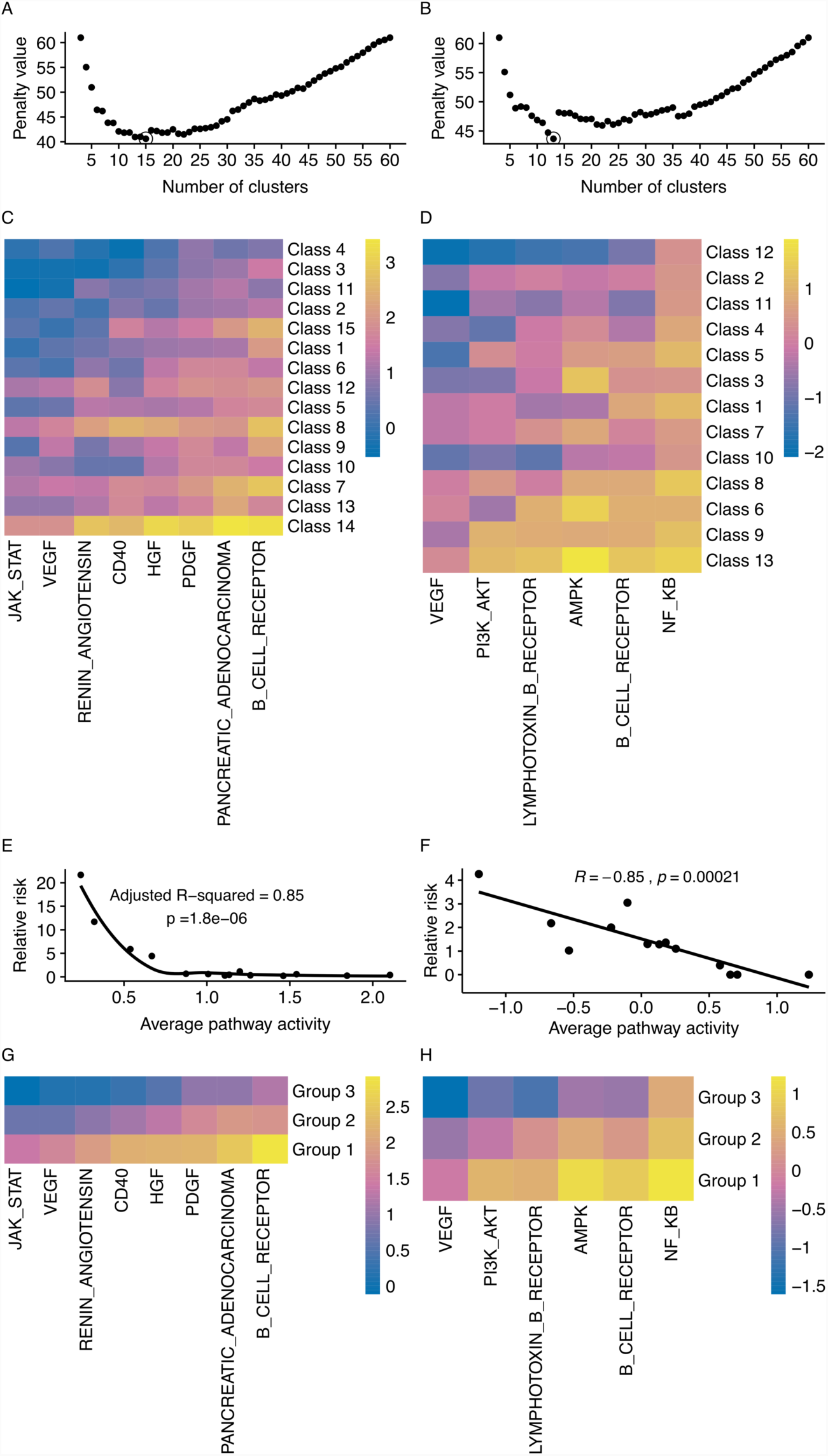
Activity spectrums of eight selected pathways allows CTRL-to-TNBC ratio-sensitive categorization of pooled Type A samples. (A) The optimum number of clusters in the eight-pathway model was determined by the penalty method for Type A. The number of clusters was selected as the lowest penalty value (circled dot). (B) The optimum number of clusters in the six-pathway model was determined by the penalty method for Type B. The number of clusters was selected as the lowest penalty value (circled dot). (C) Patterns of the eight selected pathway activity spectrums in all classes of Type A. (D) Patterns of the six selected pathway activity spectrums in all classes of Type B. (E) Correlation between the average pathway activity and the relative risk of Type A. The power law distribution curve is shown as a black line. (F) Correlation between the average pathway activity and the relative risk of Type B. The linear regression line is shown as a black line. (G) Patterns of the eight selected pathways activity spectrums in each group of Type A. (H) Patterns of the six selected pathways activity spectrums in each group of Type B.

**Table 1.** Statistics of the fifteen classes in the Type A cohort.

| **Class** | **% in**<br>**TNBC<sup>a</sup>** | **% in**<br>**CTRL<sup>b</sup>** | **Standard**<br>**deviation<sup>c</sup>** | **Z score<sup>c</sup>** | **Relative**<br>**risk** | **Group** |
| --- | --- | --- | --- | --- | --- | --- |
| 4 | 0.2115 | 0.0098 | 0.0096 | 20.9860 | 21.6663 | 3 |
| 3 | 0.1154 | 0.0099 | 0.0096 | 10.9777 | 11.7050 | 3 |
| 11 | 0.1154 | 0.0196 | 0.0136 | 7.0308 | 5.8841 | 3 |
| 2 | 0.1731 | 0.0392 | 0.0189 | 7.0801 | 4.4185 | 3 |
| 15 | 0.0769 | 0.0681 | 0.0246 | 0.3580 | 1.1292 | 2 |
| 1 | 0.0385 | 0.0591 | 0.0230 | -0.8965 | 0.6508 | 3 |
| 6 | 0.0577 | 0.0975 | 0.0293 | -1.3603 | 0.5916 | 2 |
| 12 | 0.0385 | 0.0689 | 0.0252 | -1.2095 | 0.5583 | 2 |
| 5 | 0.0385 | 0.0883 | 0.0278 | -1.7924 | 0.4357 | 2 |
| 8 | 0.0192 | 0.0488 | 0.0212 | -1.3981 | 0.3937 | 1 |
| 9 | 0.0192 | 0.0591 | 0.0231 | -1.7260 | 0.3255 | 2 |
| 10 | 0.0385 | 0.1371 | 0.0337 | -2.9298 | 0.2806 | 2 |
| 7 | 0.0192 | 0.0785 | 0.0262 | -2.2645 | 0.2451 | 1 |
| 13 | 0.0385 | 0.1770 | 0.0371 | -3.7355 | 0.2173 | 2 |
| 14 | 0 | 0.0392 | 0.0191 | -2.0518 | 0 | 1 |

<sup>a</sup>Proportions of TNBC patients in each class. <sup>b</sup>Proportions of cancer free elderly women in each class. <sup>c</sup>Standard deviations and Z scores were calculated based on random simulations.

**Table 2.** Statistics of the thirteen classes in the Type B cohort.

| **Class** | **% in**<br>**TNBC<sup>a</sup>** | **% in**<br>**CTRL<sup>b</sup>** | **Standard**<br>**deviation<sup>c</sup>** | **Z score<sup>c</sup>** | **Relative**<br>**risk** | **Group** |
| --- | --- | --- | --- | --- | --- | --- |
| 12 | 0.0926 | 0.0217 | 0.0168 | 4.2208 | 4.2608 | 3 |
| 2 | 0.1481 | 0.0486 | 0.0247 | 4.0365 | 3.0481 | 2 |
| 11 | 0.1296 | 0.0595 | 0.0272 | 2.5770 | 2.1782 | 3 |
| 4 | 0.1296 | 0.0648 | 0.0280 | 2.3132 | 1.9994 | 2 |
| 5 | 0.0741 | 0.0544 | 0.0260 | 0.7537 | 1.3605 | 2 |
| 3 | 0.0556 | 0.0429 | 0.0235 | 0.5402 | 1.2962 | 1 |
| 1 | 0.0556 | 0.0432 | 0.0231 | 0.5329 | 1.2849 | 2 |
| 7 | 0.1481 | 0.1348 | 0.0393 | 0.3387 | 1.0988 | 2 |
| 10 | 0.1111 | 0.1087 | 0.0358 | 0.0675 | 1.0223 | 3 |
| 8 | 0.0556 | 0.1407 | 0.0405 | -2.1039 | 0.3948 | 1 |
| 6 | 0 | 0.0539 | 0.0260 | -2.0739 | 0 | 2 |
| 9 | 0 | 0.1295 | 0.0387 | -3.3476 | 0 | 1 |
| 13 | 0 | 0.0971 | 0.0338 | -2.8773 | 0 | 1 |

<sup>a</sup>Proportions of TNBC patients in each class. <sup>b</sup>Proportions of cancer free elderly women in each class. <sup>c</sup>Standard deviations and Z scores were calculated based on random simulations.

Fig. 4C and Fig. 4D presented the pathway patterns in different classes that were in descending order of relative risk in Type A and Type B, respectively. The average Z-score of the pathways in each class and the relative risk followed a power law distribution (p = 1.8e-6, Fig. 4E) in Type A, while a linear correlation was found in Type B (p=2.1e-4, Fig. 4F). These finding reinforce that suppression of pathway activity is highly correlated with the germline genetics begetting TNBC vulnerability. After further analyzing the distribution statistics of each class for better interpretation, classes of Type A and those of Type B were separately merged into three groups (Table 3 for Type A and Table 4 for Type B). Compared to the classes, the groups had better-defined gradient descending patterns of the average pathway activity along with the increasing of relative risk (Fig. 4G-H), and thus provided more feasible stratifications for phenotype associating analyses.

**Table 3.** Statistics of the three groups in the Type A cohort.

| **Group** | **% in**<br>**TNBC<sup>a</sup>** | **% in**<br>**CTRL<sup>b</sup>** | **Standard**<br>**deviation<sup>c</sup>** | **Z score<sup>c</sup>** | **Relative**<br>**risk** |
| --- | --- | --- | --- | --- | --- |
| 3 | 0.6538 | 0.1366 | 0.0335 | 15.4198 | 4.7856 |
| 2 | 0.3077 | 0.6966 | 0.0450 | -8.6456 | 0.4417 |
| 1 | 0.0385 | 0.1668 | 0.0362 | -3.5422 | 0.2306 |

<sup>a</sup>Proportions of TNBC patients in each class. <sup>b</sup>Proportions of cancer free elderly women in each class. <sup>c</sup>Standard deviations and Z score were calculated based on random simulations.

**Table 4.** Statistics of the three groups in the Type B cohort.

| **Group** | **% in**<br>**TNBC<sup>a</sup>** | **% in**<br>**CTRL<sup>b</sup>** | **Standard**<br>**deviation<sup>c</sup>** | **Z score<sup>c</sup>** | **Relative**<br>**risk** |
| --- | --- | --- | --- | --- | --- |
| 3 | 0.3333 | 0.1887 | 0.0457 | 3.1624 | 1.7667 |
| 2 | 0.5556 | 0.4003 | 0.0561 | 2.7671 | 1.3879 |
| 1 | 0.1111 | 0.4110 | 0.0567 | -5.2931 | 0.2703 |

<sup>a</sup>Proportion of TNBC patients in each class. <sup>b</sup>Proportion of cancer free elderly women in each class. <sup>c</sup>Standard deviation and z score were calculated based on random simulations.

### Clinical outcomes associated with relative risk defined in groups

After further classification and annotation, Type A and Type B individuals with pathogenic or rare non-synonymous mutations in BRCA1/2 from the TNBC cohort were remarkably enriched in specific groups of distinctive relative risks. BRCA1/2 rare non-synonymous mutation carriers were enriched in Group 3 in Type A (Fig. 5A) whereas Group 2 in Type B (Fig. 5B). Based on 100,000 times of permutation of the class labels, we found the patients in class 3 (part of Group 3) of Type A were more likely to carry BRCA1/2 non-synonymous mutations with statistical significance (p = 0.0078), whereas class 1 and class 5 in Type B (both were parts of Group 2) enriched more patients carrying BRCA1/2 (p = 0.0227 and p = 0.0396, respectively) non-synonymous mutations. This divergent distribution might introduce a more accurate prediction model of the BRCA1/2 non-synonymous mutation pathogenicity when combined with population typing, specifically that BRCA1/2 non-synonymous mutation carriers in Type A have a higher risk than in Type B, which could help better assess the general disease vulnerability in individuals bearing relevant functional mutations.

**Fig. 5.**
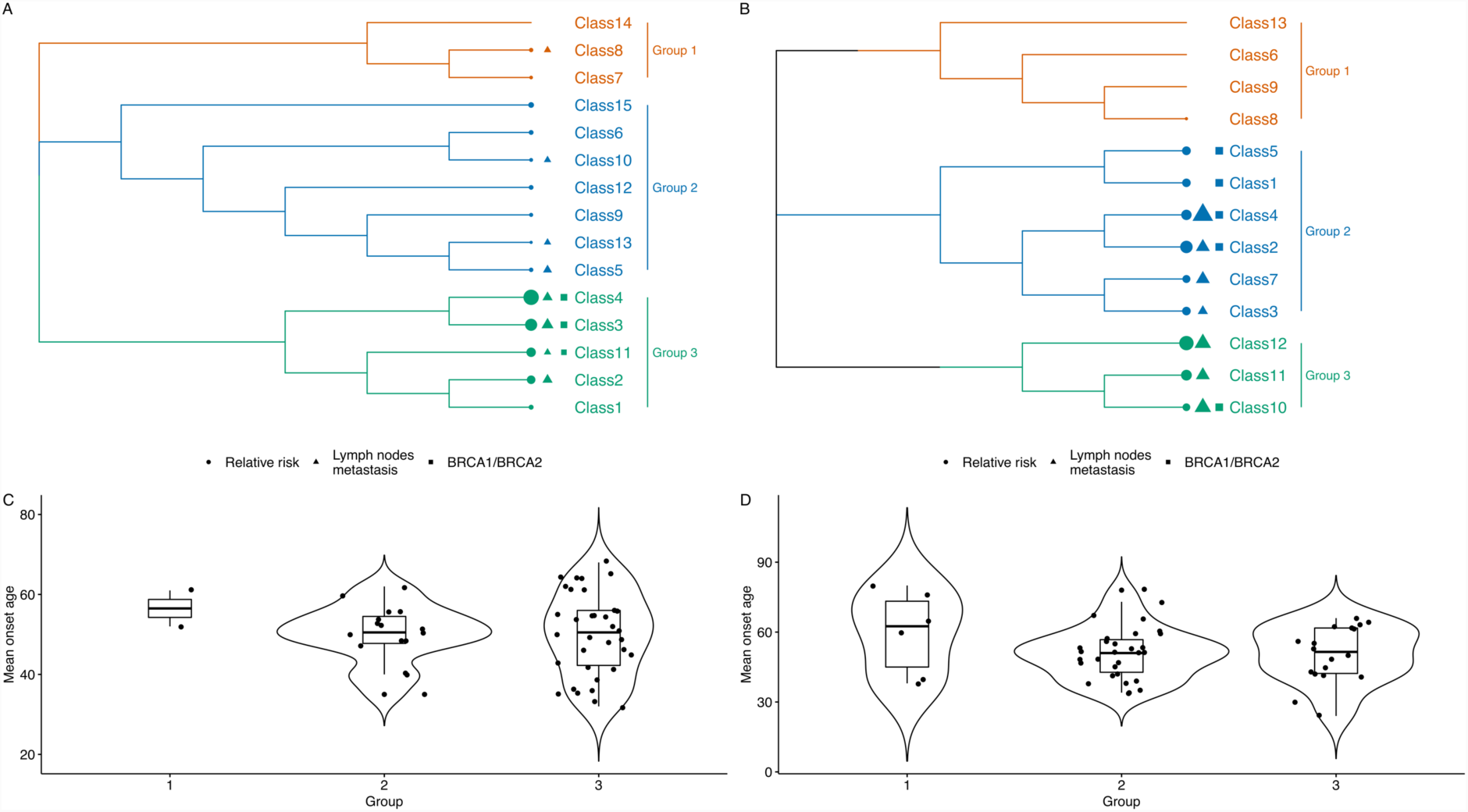
Clinical characteristics of the classes and groups. (A) Tree view of the Type A cohort showing the relative risk of each class along with the distribution of lymph nodes metastasis and BRCA1/2 mutations. (B) Tree view of the Type B cohort showing the relative risk of each class along with the distribution of lymph nodes metastasis and BRCA1/2 mutations. The size of the circles, triangles, and squares represent the distributions, which were only comparable within types but not across types. (C) Violin plot of the onset age for the three groups in Type A. (D) Violin plot of the onset age for the three groups in Type B.

Lymph node metastasis and age of onset could also be associated with the relative risk calculated by this model. Statistical annotation showed that lymph node metastasis was associated with class 3 (p = 0.0014, relative risk = 11.71), class 5 (p < 0.0001, relative risk = 0.44), class 8 (p < 0.0001, relative risk = 0.39) in Type A, as well as class 4 (p = 0.0038, relative risk = 2.0) and 12 (p = 0.0428, relative risk = 4.26) in Type B (Fig. 5A-B). With respect to the age of onset of TNBC, a gradual decrease was observed along with the increasing relative risk from Group 1 to Group 3 in both Type A and Type B (Fig. 5C-D), where patients in Group 1 of Type B showed a significantly higher onset age (p = 0.029, relative risk = 0.27).

## Discussion

Previous screening approaches for genetic predisposition were generally based on partial high-penetrance gene mutations. These approaches found only a limited number of breast cancer cases were attributable to heredofamilial disorders (52), leaving the vast majority of cases as sporadic-like, including TNBC (18). We hypothesized that these sporadic-like cases involved germline mutations. Such sporadic-like cases often display remarkable non-Mendelian genetic features, such as polygenic traits, incomplete- and co-dominance (53,54). On the one hand, these features make it difficult to rule out familial inheritance, but on the other hand favor our hypothesis of using holistic germline information to characterize sporadic-like TNBC. Our approach assumes that rare non-synonymous germline mutations predispose individuals in a population to cancer, whereas somatic mutations initiated and triggered the progression of the cancer (55). Furthermore, we postulated that breast cancer is a ‘bubbling’ and persistent pathogenic stress on the female population, and that resistance against this stress is evolutionary selected. As an evolutionary population trait, the resistance must be encoded in a complex multi-factorial genetic network, which can only be disrupted by the collective effects of multiple mutational events. These assumptions underlie our rationale to integrate the holistic information of all rare mutations in the entire coding regions of germline DNA to evaluate the quantitative status of this resistance.

Our knowledge-data-driven hybrid machine learning framework can project layer by layer high-dimensional (more than ten thousand), inscrutable, binary, and sparse mutation data onto lower-dimensional (dozens) but information-rich data with quantitative features. The data is integrated and transformed by the built-in machine learning algorithms, and each layer can take the output of the previous layer as the input to generate the next output. We use this approach to evaluate the risk of developing TNBC in an individual. The model “comprehends” the rare genetic mutations in each female as a whole, assesses the level of risk and classifies them as having either high or low risk accordingly. Using our machine learning approach to understand the overall genetic information, we found that in the Type A population, TNBC patients could be separated from the CTRL individuals (AUC = 0.89, Fig. 2D). This finding led to the first key conclusion that in patients with sporadic TNBC, the cause of disease might be systematically rooted in the germline mutations. In addition, the driving force of the etiology in Type A and Type B cohorts followed in an analogous manner, as seen with an r of 0.70 (p = 4.5e-10, Fig. 3E) in the significance correlation analysis.

More specifically, immune-related pathways, such as B-cell receptor, NF-κB, CD40, and lymphotoxin-B receptor signaling, were found to be involved in the relationship between pathways activities and TNBC risk. The degree of suppression in these pathways was rigorously correlated with the relative risk for TNBC when compared with CTRL. The highest levels of suppression were significantly associated with increased risk in both Type A and Type B. Furthermore, pooled Type A and Type B cohorts were clustered into three distinct groups in accordance to their pathway activity spectrum. The relative risk in the TNBC cases was calculated compared with the CTRL within each of the three groups. The relative risk intrinsically indicates the possible risk in any randomly selected sample of CTRL or TNBC in any given group with similar pathway activity spectra, and thus quantitatively describes the TNBC risk in the given activity group. We observed the relative risk increased from group 1 to group 3 in both Type A and Type B, whereas the onset age gradually decreased. The lymph node metastasis was mainly found in group 2 and group 3 in both Type A and Type B. Patients with BRCA1/2 mutations were more likely to be found in group 3 of Type A and in group 2 of Type B.

We used a cohort of cancer-free elderly women as the control. This group had an average age above 80, which we believe represents the minimum risk under specific ethnic and geographical conditions, as they were free of cancer for these many years. This enabled us to map the TNBC risk according to the relative risk of each pathway activity group. However, this sample of healthy senior females might not fully represent general ethnic and geographical populations, as well as age ranges. More representative samples will be needed in future studies to link other phenotypical traits with the pathway activity spectra through our model.

## Conclusions

Our machine learning framework (DAGG) views the germline information buried in individual genomes differently. This model does not consider for individual mutation separately in the gene level but projects all rare germline mutations into each of the signaling pathways in the functional level (signaling pathway activities) and then uses the spectra that consist of multiple pathways to delineate pathological characters of each individual. The results have validated the hypothesis that relative risks indicated by the germline-decided pathway activity in DAGG could serve as an early index for predicting TNBC related pathology. The TNBC related pathology could include but not limited to onset, metastasis relevant prognosis and resistance against pathogenic mutations. These provides us two advantages: the first is the possibility of early identification of TNBC high-risk. This allows better treatment and improves prognosis. The second is the methodological generality of DAGG that enables applications in other type of genetic-related cancer types once training datasets are properly selected.

## Acknowledgement

The authors would like to thank Yanxiang Ni, Jin Gu, Xu Li, and Zhonghai Zhang for feedback on the manuscript and helpful discussions.

## Author contributions

GN and YF conceived the experiments. GN, KW, MY, and ZW, designed the experiments. MY, QZ, XL, TZ, MC, JX, CY, and HG performed the experiments. YF, GN, QZ, SH, CZ, GT, YQS, and MQZ analyzed the data. YF, MY, ZF, and GN wrote the paper. All authors discussed the results and contributed to the final manuscript.

## References

1. Chen W, Zheng R, Baade PD, Zhang S, Zeng H, Bray F, et al. Cancer statistics in China, 2015. CA Cancer J Clin 2016;66:115–32

2. Jiang X, Tang H, Chen T. Epidemiology of gynecologic cancers in China. J Gynecol Oncol 2018;29:e7

3. Siegel RL, Miller KD, Jemal A. Cancer statistics, 2018. CA Cancer J Clin 2018;68:7–30

4. Reis-Filho JS, Tutt AN. Triple negative tumours: a critical review. Histopathology 2008;52:108–18

5. Couch FJ, Hart SN, Sharma P, Toland AE, Wang X, Miron P, et al. Inherited mutations in 17 breast cancer susceptibility genes among a large triple-negative breast cancer cohort unselected for family history of breast cancer. J Clin Oncol 2015;33:304–11

6. Dent R, Trudeau M, Pritchard KI, Hanna WM, Kahn HK, Sawka CA, et al. Triple-negative breast cancer: clinical features and patterns of recurrence. Clin Cancer Res 2007;13:4429–34

7. Gonzalez-Angulo AM, Timms KM, Liu S, Chen H, Litton JK, Potter J, et al. Incidence and outcome of BRCA mutations in unselected patients with triple receptor-negative breast cancer. Clin Cancer Res 2011;17:1082–9

8. Hartman AR, Kaldate RR, Sailer LM, Painter L, Grier CE, Endsley RR, et al. Prevalence of BRCA mutations in an unselected population of triple-negative breast cancer. Cancer 2012;118:2787–95

9. Sharma P, Klemp JR, Kimler BF, Mahnken JD, Geier LJ, Khan QJ, et al. Germline BRCA mutation evaluation in a prospective triple-negative breast cancer registry: implications for hereditary breast and/or ovarian cancer syndrome testing. Breast Cancer Res Treat 2014;145:707–14

10. Wong-Brown MW, Meldrum CJ, Carpenter JE, Clarke CL, Narod SA, Jakubowska A, et al. Prevalence of BRCA1 and BRCA2 germline mutations in patients with triple-negative breast cancer. Breast Cancer Res Treat 2015;150:71–80

11. Giordano TJ. The cancer genome atlas research network: a sight to behold. Endocr Pathol 2014;25:362–5

12. Hahnen E, Hauke J, Engel C, Neidhardt G, Rhiem K, Schmutzler RK. Germline Mutations in Triple-Negative Breast Cancer. Breast Care 2017;12:15–9

13. Brady-West DC, McGrowder DA. Triple negative breast cancer: therapeutic and prognostic implications. Asian Pac J Cancer Prev 2011;12:2139–43

14. Bianchini G, Balko JM, Mayer IA, Sanders ME, Gianni L. Triple-negative breast cancer: challenges and opportunities of a heterogeneous disease. Nat Rev Clin Oncol 2016;13:674–90

15. Malorni L, Shetty PB, De Angelis C, Hilsenbeck S, Rimawi MF, Elledge R, et al. Clinical and biologic features of triple-negative breast cancers in a large cohort of patients with long-term follow-up. Breast Cancer Res Treat 2012;136:795–804

16. Denkert C, Liedtke C, Tutt A, von Minckwitz G. Molecular alterations in triple-negative breast cancer-the road to new treatment strategies. Lancet 2017;389:2430–42

17. Lehmann BD, Bauer JA, Chen X, Sanders ME, Chakravarthy AB, Shyr Y, et al. Identification of human triple-negative breast cancer subtypes and preclinical models for selection of targeted therapies. J Clin Invest 2011;121:2750–67

18. Lobo M, Lopez-Tarruella S, Luque S, Lizarraga S, Flores-Sanchez C, Bueno O, et al. Evaluation of Breast Cancer Patients with Genetic Risk in a University Hospital: Before and After the Implementation of a Heredofamilial Cancer Unit. J Genet Couns 2017

19. Plevritis SK, Munoz D, Kurian AW, Stout NK, Alagoz O, Near AM, et al. Association of Screening and Treatment With Breast Cancer Mortality by Molecular Subtype in US Women, 2000-2012. JAMA 2018;319:154–64

20. Anderson K, Thompson PA, Wertheim BC, Martin L, Komenaka IK, Bondy M, et al. Family history of breast and ovarian cancer and triple negative subtype in hispanic/latina women. Springerplus 2014;3:727

21. von Minckwitz G, Schneeweiss A, Loibl S, Salat C, Denkert C, Rezai M, et al. Neoadjuvant carboplatin in patients with triple-negative and HER2-positive early breast cancer (GeparSixto; GBG 66): a randomised phase 2 trial. Lancet Oncol 2014;15:747–56

22. Stevens KN, Vachon CM, Couch FJ. Genetic susceptibility to triple-negative breast cancer. Cancer Res 2013;73:2025–30

23. Easton DF, Pharoah PD, Antoniou AC, Tischkowitz M, Tavtigian SV, Nathanson KL, et al. Gene-panel sequencing and the prediction of breast-cancer risk. N Engl J Med 2015;372:2243–57

24. Kiiski JI, Pelttari LM, Khan S, Freysteinsdottir ES, Reynisdottir I, Hart SN, et al. Exome sequencing identifies FANCM as a susceptibility gene for triple-negative breast cancer. Proc Natl Acad Sci U S A 2014;111:15172–7

25. Meindl A, Hellebrand H, Wiek C, Erven V, Wappenschmidt B, Niederacher D, et al. Germline mutations in breast and ovarian cancer pedigrees establish RAD51C as a human cancer susceptibility gene. Nat Genet 2010;42:410–4

26. Neidhardt G, Hauke J, Ramser J, Gross E, Gehrig A, Muller CR, et al. Association Between Loss-of-Function Mutations Within the FANCM Gene and Early-Onset Familial Breast Cancer. JAMA Oncol 2017;3:1245–8

27. Ramus SJ, Song H, Dicks E, Tyrer JP, Rosenthal AN, Intermaggio MP, et al. Germline Mutations in the BRIP1, BARD1, PALB2, and NBN Genes in Women With Ovarian Cancer. J Natl Cancer Inst 2015;107

28. Walsh CS. Two decades beyond BRCA1/2: Homologous recombination, hereditary cancer risk and a target for ovarian cancer therapy. Gynecol Oncol 2015;137:343–50

29. Gonzalez-Rivera M, Lobo M, Lopez-Tarruella S, Jerez Y, Del Monte-Millan M, Massarrah T, et al. Frequency of germline DNA genetic findings in an unselected prospective cohort of triple-negative breast cancer patients participating in a platinum-based neoadjuvant chemotherapy trial. Breast Cancer Res Treat 2016;156:507–15

30. von Minckwitz G, Loibl S, Untch M, Eidtmann H, Rezai M, Fasching PA, et al. Survival after neoadjuvant chemotherapy with or without bevacizumab or everolimus for HER2-negative primary breast cancer (GBG 44-GeparQuinto)dagger. Ann Oncol 2014;25:2363–72

31. von Minckwitz G, Rezai M, Fasching PA, Huober J, Tesch H, Bauerfeind I, et al. Survival after adding capecitabine and trastuzumab to neoadjuvant anthracycline-taxane-based chemotherapy for primary breast cancer (GBG 40--GeparQuattro). Ann Oncol 2014;25:81–9

32. Zhang J, Sun J, Chen J, Yao L, Ouyang T, Li J, et al. Comprehensive analysis of BRCA1 and BRCA2 germline mutations in a large cohort of 5931 Chinese women with breast cancer. Breast Cancer Res Treat 2016;158:455–62

33. Kotsopoulos J, Lubinski J, Gronwald J, Cybulski C, Demsky R, Neuhausen SL, et al. Factors influencing ovulation and the risk of ovarian cancer in BRCA1 and BRCA2 mutation carriers. Int J Cancer 2015;137:1136–46

34. Audeh MW, Carmichael J, Penson RT, Friedlander M, Powell B, Bell-McGuinn KM, et al. Oral poly(ADP-ribose) polymerase inhibitor olaparib in patients with BRCA1 or BRCA2 mutations and recurrent ovarian cancer: a proof-of-concept trial. Lancet 2010;376:245–51

35. Byrski T, Huzarski T, Dent R, Marczyk E, Jasiowka M, Gronwald J, et al. Pathologic complete response to neoadjuvant cisplatin in BRCA1-positive breast cancer patients. Breast Cancer Res Treat 2014;147:401–5

36. Ledermann J, Harter P, Gourley C, Friedlander M, Vergote I, Rustin G, et al. Olaparib maintenance therapy in patients with platinum-sensitive relapsed serous ovarian cancer: a preplanned retrospective analysis of outcomes by BRCA status in a randomised phase 2 trial. Lancet Oncol 2014;15:852–61

37. Pennington KP, Walsh T, Harrell MI, Lee MK, Pennil CC, Rendi MH, et al. Germline and somatic mutations in homologous recombination genes predict platinum response and survival in ovarian, fallopian tube, and peritoneal carcinomas. Clin Cancer Res 2014;20:764–75

38. Tutt A, Robson M, Garber JE, Domchek SM, Audeh MW, Weitzel JN, et al. Oral poly(ADP-ribose) polymerase inhibitor olaparib in patients with BRCA1 or BRCA2 mutations and advanced breast cancer: a proof-of-concept trial. Lancet 2010;376:235–44

39. Brunello A, Borgato L, Basso U, Lumachi F, Zagonel V. Targeted approaches to triple-negative breast cancer: current practice and future directions. Curr Med Chem 2013;20:605–12

40. Khera AV, Chaffin M, Aragam KG, Haas ME, Roselli C, Choi SH, et al. Genome-wide polygenic scores for common diseases identify individuals with risk equivalent to monogenic mutations. Nature Genetics 2018

41. Sundaram L, Gao H, Padigepati SR, McRae JF, Li Y, Kosmicki JA, et al. Predicting the clinical impact of human mutation with deep neural networks. Nature Genetics 2018

42. Zhou J, Theesfeld CL, Yao K, Chen KM, Wong AK, Troyanskaya OG. Deep learning sequence-based ab initio prediction of variant effects on expression and disease risk. Nat Genet 2018

43. Wang B, Bao S, Zhang Z, Zhou X, Wang J, Fan Y, et al. A rare variant in MLKL confers susceptibility to ApoE ε4-negative Alzheimer’s disease in Hong Kong Chinese population. Neurobiol Aging 2018;68:160.e1–.e7

44. Stovgaard ES, Nielsen D, Hogdall E, Balslev E. Triple negative breast cancer – prognostic role of immune-related factors: a systematic review. Acta Oncologica 2018;57:74–82

45. Li H. Aligning sequence reads, clone sequences and assembly contigs with BWA-MEM. q-bioGN 2013;arXiv:1303.3997v2

46. McKenna AH, Hanna M, Banks E, Sivachenko A, Cibulskis K, Kernytsky A, et al. The Genome Analysis Toolkit: A MapReduce framework for analyzing next-generation DNA sequencing data. Genome Res 2010;20:1297–303

47. Wang K, Li M, Hakonarson H. ANNOVAR: functional annotation of genetic variants from high-throughput sequencing data. Nucleic Acids Res 2010;38:e164–e

48. Murtagh F, Legendre P. Ward’s Hierarchical Agglomerative Clustering Method: Which Algorithms Implement Ward’s Criterion? J Classif 2014;31:274–95

49. Kelley LA, Gardner SP, Sutcliffe MJ. An automated approach for clustering an ensemble of NMR-derived protein structures into conformationally related subfamilies. Protein Eng Des Sel 1996;9:1063–5

50. Iorio F, Knijnenburg Theo A, Vis Daniel J, Bignell Graham R, Menden Michael P, Schubert M, et al. A Landscape of Pharmacogenomic Interactions in Cancer. Cell 2016;166:740–54

51. Kelley LA, Gardner SP, Sutcliffe MJ. An automated approach for clustering an ensemble of NMR-derived protein structures into conformationally related subfamilies. Protein Eng Des Sel 1996;9:1063–5

52. Berry DA, Cronin KA, Plevritis SK, Fryback DG, Clarke L, Zelen M, et al. Effect of screening and adjuvant therapy on mortality from breast cancer. N Engl J Med 2005;353:1784–92

53. Antoniou AC, Easton DF. Models of genetic susceptibility to breast cancer. Oncogene 2006;25:5898–905

54. Jara L, Morales S, de Mayo T, Gonzalez-Hormazabal P, Carrasco V, Godoy R. Mutations in BRCA1, BRCA2 and other breast and ovarian cancer susceptibility genes in Central and South American populations. Biol Res 2017;50:35

55. Low SK, Zembutsu H, Nakamura Y. Breast cancer: The translation of big genomic data to cancer precision medicine. Cancer Sci 2017;109:497–506

