## Supplementary Tables for "Rare non-synonymous germline mutations systematically define the risk of triple negative breast cancer"

**Supplementary Table 1 Patients and Disease Characteristics.**

| Characteristic | Baseline | Post-surgry |
| --- | --- | --- |
| Age, Mean (range，years) | 51(24-78) | - |
| Histologic type, n(%) |  |  |
| Ductal | 62/66 (93.9) | 38/40(95.0) |
| Other | 4/66(6.1) | 2/40(5.0) |
| Histological grade, n (%) |  |  |
| 1 | 0/66 (0.0) | 0/40(0.0) |
| 2 | 20/66 (30.3) | 13/40(32.5) |
| 3 | 41/66 (62.1) | 22/40(55.0) |
| Unknown | 5/66 (7.6) | 5/40(12.5) |
| Tumor size, n (%) |  |  |
| ≤2cm | 17/66(25.8) | 11/40 (27.5) |
| ＞2cm | 45/66 (68.2) | 29/40 (72.5) |
| Unknown | 4/66 (6.1) | 0/40 (0.0) |
| Nodal status, n (%) |  |  |
| LN- | 39/66 (59.1) | 28/40 (70.0) |
| LN+ | 23/66 (34.8) | 12/40 (30.0) |
| Unknown | 4/66 (6.1) | 0/40 (0.0) |
| Ki 67 status, n (%) |  |  |
| ≤10 | 5/66 (7.6) | 3/40(7.5) |
| 10-30  ＞30  unknown | 11/66 (16.7)  41/66 (62.1)  9/66(13.6) | 4/40(10.0)  32/40(80.0)  1/40(2.5) |

**Supplementary Table 2 ER, PR, and HER2 status and TNM staging of the TNBC patients.**

| ID | Age | lesion site^*^ | T | N | ER | PR | CerbB2 | FISH(Her2) | Ki67 |
| --- | --- | --- | --- | --- | --- | --- | --- | --- | --- |
| ZY101 | 34 | 1 | 2 | 0 | 0, <1% | 0, <1% | 1+ | - | 40% |
| ZY102 | 38 | 1 | 3 | 1 | 1+, <1% | 0 | 0 |  | 50% |
| ZY103 | 63 | 1 | 2 | 1 | 0 | 1+, 2% | 2+ | - | 40% |
| ZY104 | 53 | 1 | 2 | 1 | 1+, <1% | 1+, <1% | 2+ | - | 5% |
| ZY105 | 55 | 3 | 2 | 1 | 1+, <1% | 0, <1% | 1+ |  | 75% |
| ZY106 | 56 | 1 | 2 | 0 | 0 | 0 | 0 | - | 85% |
| ZY107 | 64 | 2 | 3 | 1 | 1+, <1% | 1+, 2% | 2+ | - | 45% |
| ZY108 | 53 | 2 | 1c | 0 | 1+, <1% | 1+, <1% | 0 |  | 25% |
| ZY109 | 57 | 1 | 2 | 1 | 0 | 0 | 2+ | - | 70% |
| ZY110 | 49 | 1 | 2 | 0 | 0 | 0 | 0 |  | 30% |
| ZY111 | 40 | 1 | 2 | 0 | 0 | 0 | 0 | - | 30% |
| ZY112 | 65 | 2 | 1b | 0 | 0 | 0 | 0 | - | 70% |
| ZY113 | 60 | 2 | 2 | 0 | 1+, <1% | 1+, <1% | 0 | - | 55% |
| ZY114 | 30 | 2 | 2 | 0 | 1+, 1% | 1+, <1% | 0 | - | 80% |
| ZY115 | 65 | 2 | 2 | 0 | 1+, <1% | 1+, <1% | 2+ | - | 35% |
| ZY116 | 64 | 1 | 4 | 0 | 0, <1％ | 0, <1％ | 1+ | - | 85% |
| ZY117 | 68 | 2 | 1c | 1 | 1+, <1% | 1+, <1% | 1+ | - | 70% |
| ZY118 | 61 | 1 | 2 | 1 | 0 | 1+, 10% | 0 |  | 90% |
| ZY119 | 52 | 1 | 1c | 0 | 1+, <1% | 1+, <1% | 2+ | - | 60% |
| ZY120 | 73 | 2 | 1c | 1 | 0 | 1+, 30% | 0 |  | 80% |
| ZY121 | 35 | 2 | 2 | 2 | 2+, 2% | 2+, 3% | 0 | - | 85%+ |
| ZY122 | 45 | 2 | 1c | 1 | 0 | 1+, 2% | 0 | - | 10% |
| ZY123 | 66 | 2 | 2 | 0 | 0 | 0 | 2+ | - | 40% |
| ZY124 | 56 | 2 | 2 | 1 | 1+, <1% | 1+, <1% | 1+ | - | 40% |
| ZY125 | 46 | 1 | 3 | 3 | 0 | 1% | 0 |  | 80% |
| ZY126 | 24 | 1 | 2 | 0 | 1+, <1% | 1+, <1% | 1+ | - | 80% |
| ZY127 | 50 | 2 | 2 | 0 | 1+, <1% | 1+, <1% | 0 |  | 80% |
| ZY128 | 49 | 1 | 2 | 0 | 1+, <1% | 1+, <1% | 1+ | - | 50% |
| ZY129 | 62 | 1 | 1c | 0 | 0 | 1+, <1% | 0 | - | 40% |
| ZY130 | 48 | 2 | 2 | 1 | 1+, <1% | 1+, <1% | 0 |  | 70%+ |
| ZY131 | 62 | 2 | 4 | 2 | 0, <1% | 0, <1% | 0 |  | 30% |
| ZY132 | 42 | 1 | 2 | 0 | 0 | 0 | 1+ | - | 50% |
| ZY133 | 39 | 1 | 2 | 0 | 0 | 0 | 0 |  | 20% |
| ZY134 | 55 | 2 | 1c | 0 | 0, <1% | 0, <1% | 1+ |  | 40% |
| ZY135 | 47 | 1 | 2 | 0 | 0 | 0 | 0 | - | 60% |
| ZY136 | 51 | 1 | 2 | 1 | 0 | 0 | 0 |  | 65% |
| ZY137 | 41 | 1 | 2 | 0 | 0 | 0 | 0 | - | 20% |
| ZY138 | 38 | 1 | 2 | 0 | 1+, <1% | 2+, 2% | 1+ | - | 70% |
| ZY139 | 32 | 1 | 1 | 0 | 0 | 0 | 0 | 1+ | 25% |
| ZY140 | 53 | 1 | 2 | 0 | <1% | 0 | 2+ | - | 15% |
| ZY141 | 41 | 1 | 2 | 0 | 1+, <1% | 1+ ＜1% | 0 |  | 20% |
| ZY142 | 41 | 2 | 2 | 1 | 0, <1% | 0, <1% | 2+ | - | 30% |
| ZY143 | 36 | 1 | 2 | 0 | 1+, 1% | 1+, 1% | 0 | - | 30% |
| ZY144 | 50 | 1 | 2 | 1 | 0 | 0 | 0 | - | 50% |
| ZY145 | 38 | 1 | 1 | 0 | 0 | 0 | 1+ | - | 20% |
| ZY146 | 58 | 2 | 1 | 0 | 0 | 0 | 1+ | - | 50% |
| ZY147 | 33 | 1 | 1c | 0 | 0 | 0 | 2+ | - | 70% |
| ZY148 | 52 | 2 | 1 | 0 | 1+, <1% | 1+, <1% | 1+ | - | 60% |
| ZY149 | 60 |  | 1C | 1 | 0 | 2+, 50% | 0 |  | 70% |
| ZY150 | 60 | 1 | 1 | 0 | 1+, <1% | 1+,＜1% | 2+ | - | 50% |
| ZY151 | 54 |  | 1 | 0 | 0 | 0 | 0 | 0 | 50% |
| ZY152 | 53 | 1 | 2 | 1 | 0 | 1+, <1% | 1+ | - | 40% |
| ZY153 | 62 | 2 | 2 | 1 | - | 1+,＜5% | - | - | 10% |
| ZY154 | 61 | 2 | 2 | 1 | - | - | - | - | 10% |
| ZY155 | 40 | 1 | 2 | 0 | 1+, <1% | 1+, 5% | 0 |  | 40% |
| ZY156 | 41 | 1 | 2(2.6*2.3cm) | 0 | 0 | 0 | 0 |  | 70% |
| ZY157 | 47 | 2 | 1c | 0 | 0 | 0 | 1+ | - | 90% |
| ZY158 | 54 | 2 | 1c | 0 | 1+, <1% | 1+, <1% | 0 | - | 50% |
| ZY159 | 80 | 2 | 1c | 0 | 0 | 0 | 0 | - | 5% |
| ZY160 | 42 | 1 | 1c(1.2*0.6cm) | 0 | 1+, <1% | 1+, <1% | 2+ | - | 90% |
| ZY161 | 36 | 2 | 2 | 0 | 0 | 0 | 0 | 0 | 40% |
| ZY162 | 53 | 2 | 2 | 0 | 0 | 0 | 2+ | - | 40% |
| ZY163 | 56 | 1 | 2 | 1 | 1+, <1% | 1+, <1% | - | - | 60% |
| ZY164 | 47 | 2 | 2 | 0 | 0 | 0 | 0 | 0 | 30% |
| ZY165 | 40 | 2 | 2 | 0 | 0 | 0 | 2+ | - | 40% |
| ZY166 | 51 | 1 | 2 | 0 | 0 | 0 | 0 |  | 80% |
| ZY167 | 56 | 2 | 2 | 0 | 0 | 0 | 0 |  | 90% |
| ZY168 | 48 | 1 | 2 | 1 | 0 | 0 | 0 |  | 80% |
| ZY169 | 50 | 2 | 1 | 0 | 0 | 0 | 0 |  | 90% |
| ZY170 | 48 | 1 | 3 | 0 | 0 | 0 | 0 |  | 90% |
| ZY171 | 48 | 1 | 2 | 0 | 0 | 0 | 0 |  | 80% |
| ZY172 | 52 | 2 | 2 | 0 | 0 | 0 | 0 |  | 90% |
| ZY173 | 62 | 2 | 1 | 2 | 0 | 0 | 0 |  | 60% |
| ZY174 | 56 | 2 | 2 | 1 | 0 | 0 | 0 |  | 40% |
| ZY175 | 59 | 1 | 2 | 0 | 0 | 0 | 0 |  | 90% |
| ZY176 | 48 | 1 | 1 | 1 | 0 | 0 | 0 |  | 90% |
| ZY177 | 43 | 1 | 2 | 0 | 0 | 0 | 0 |  | 70% |
| ZY178 | 48 | 1 | 2 | 0 | 0 | 0 | 0 |  | 80% |
| ZY179 | 43 | 1 | 2 | 0 | 0 | 0 | 0 |  | 80% |
| ZY180 | 52 | 1 | 2 | 0 | 0 | 0 | 0 |  | 25% |
| ZY181 | 39 | 2 | 2 | 0 | 0 | 0 | 0 |  | 50% |
| ZY182 | 61 | 1 | 2 | 2 | 0 | 0 | 0 |  | 90% |
| ZY183 | 76 | 1 | 1 | 0 | 0 | 0 | 1+ |  | 90% |
| ZY184 | 55 | 2 | 2 | 2 | 0 | 0 | 0 |  | 70% |
| ZY185 | 45 | 1 | 2 | 0 | 0 | 0 | 0 |  | 60% |
| ZY186 | 35 | 1 | 1 | 0 | 0 | 0 | 0 |  | 90% |
| ZY187 | 59 | 2 | 2 | 2 | 0 | 0 | 0 |  | 20% |
| ZY188 | 35 | 2 | 1 | 0 | 0 | 0 | 0 |  | 50% |
| ZY189 | 56 | 2 | 2 | 0 | 0 | 0 | 0 |  | 60% |
| ZY190 | 45 | 2 | 2 | 0 | 0 | 0 | 0 |  | 60% |
| ZY191 | 51 | 2 | 2 | 0 | 0 | 0 | 0 |  | 95% |
| ZY192 | 78 | 1 | 1 | 0 | 0 | 0 | 0 |  | 5% |
| ZY193 | 34 | 1 | 2 | 1 | 0 | 0 | 0 |  | 90% |
| ZY194 | 64 | 2 | 1 | 0 | 0 | 0 | 0 |  | 50% |
| ZY195 | 35 | 1 | 2 | 0 | 0 | 0 | 0 |  | 90% |
| ZY196 | 51 | 1 | 2 | 0 | 0 | 0 | 0 |  | 90% |
| ZY197 | 42 | 1 | 1 | 0 | 0 | 0 | 0 |  | 80% |
| ZY198 | 64 | 1 | 2 | 1 | 0 | 0 | 0 |  | 70% |
| ZY199 | 66 | 1 | 1 | 0 | 0 | 0 | 0 |  | 50% |
| ZY200 | 51 | 2 | 2 | 1 | 0 | 0 | 0 |  | 90% |
| ZY201 | 46 | 2 | 2 | 0 | 0 | 0 | 0 |  | 90% |
| ZY202 | 67 | 1 | 2 | 0 | 0 | 0 | 0 |  | 30% |
| ZY203 | 55 | 2 | 2 | 0 | 0 | 0 | 0 |  | 90% |
| ZY204 | 50 | 2 | is | 0 | 0 | 0 | 0 |  | 20% |
| ZY205 | 61 | 1 | 2 | 1 | 0 | 0 | 0 |  | 5% |
| ZY206 | 54 | 1 | 2 | 1 | 0 | 0 | 0 |  | 95% |

*lesion site: 1-left, 2-right, 3-bilateral. T: tumor size. N: regional lymph node metastasis. ER: estrogen receptor. PR: progesterone receptor.

**Supplementary Table 3 Age, sex, BRCA1/2 status, and lymph nodes metastasis status of TNBC patients and their distribution in each type, class, and group.**

| Index | Type | Class | Group | Age | Sex | BRCA1/  BRCA2^a^ | Lymph nodes  metastasis^b^ |
| --- | --- | --- | --- | --- | --- | --- | --- |
| 1 | A | 7 | 1 | 52 | female | 0 | 0 |
| 2 | A | 8 | 1 | 61 | female | 0 | 1 |
| 3 | A | 5 | 2 | 54 | female | 0 | 1 |
| 4 | A | 5 | 2 | 62 | female | 0 | 1 |
| 5 | A | 6 | 2 | 56 | female | 0 | 0 |
| 6 | A | 6 | 2 | 50 | female | 0 | 0 |
| 7 | A | 6 | 2 | 60 | female | 0 | 0 |
| 8 | A | 9 | 2 | 53 | female | 0 | 0 |
| 9 | A | 10 | 2 | 52 | female | 0 | 0 |
| 10 | A | 10 | 2 | 50 | female | 0 | 1 |
| 11 | A | 12 | 2 | 40 | female | 0 | 0 |
| 12 | A | 12 | 2 | 40 | female | 0 | 0 |
| 13 | A | 13 | 2 | 48 | female | 0 | 0 |
| 14 | A | 13 | 2 | 51 | female | 0 | 1 |
| 15 | A | 15 | 2 | 56 | female | 0 | 0 |
| 16 | A | 15 | 2 | 48 | female | 0 | 0 |
| 17 | A | 15 | 2 | 35 | female | 0 | 0 |
| 18 | A | 15 | 2 | 47 | female | 0 | 0 |
| 19 | A | 1 | 3 | 49 | female | 0 | 0 |
| 20 | A | 1 | 3 | 54 | female | 0 | 0 |
| 21 | A | 2 | 3 | 61 | female | 0 | 2 |
| 22 | A | 2 | 3 | 55 | female | 0 | 0 |
| 23 | A | 2 | 3 | 61 | female | 0 | 1 |
| 24 | A | 2 | 3 | 50 | female | 0 | 0 |
| 25 | A | 2 | 3 | 49 | female | 0 | 0 |
| 26 | A | 2 | 3 | 33 | female | 0 | 0 |
| 27 | A | 2 | 3 | 41 | female | 0 | 0 |
| 28 | A | 2 | 3 | 62 | female | 0 | 2 |
| 29 | A | 2 | 3 | 52 | female | 0 | 0 |
| 30 | A | 3 | 3 | 56 | female | 1 | 1 |
| 31 | A | 3 | 3 | 68 | female | 0 | 1 |
| 32 | A | 3 | 3 | 46 | female | 0 | 3 |
| 33 | A | 3 | 3 | 36 | female | 0 | 0 |
| 34 | A | 3 | 3 | 48 | female | 0 | 1 |
| 35 | A | 3 | 3 | 35 | female | 2 | 2 |
| 36 | A | 4 | 3 | 43 | female | 0 | 0 |
| 37 | A | 4 | 3 | 39 | female | 0 | 0 |
| 38 | A | 4 | 3 | 55 | female | 0 | 2 |
| 39 | A | 4 | 3 | 35 | female | 0 | 0 |
| 40 | A | 4 | 3 | 64 | female | 2 | 0 |
| 41 | A | 4 | 3 | 64 | female | 0 | 0 |
| 42 | A | 4 | 3 | 56 | female | 0 | 1 |
| 43 | A | 4 | 3 | 36 | female | 0 | 0 |
| 44 | A | 4 | 3 | 42 | female | 0 | NA |
| 45 | A | 4 | 3 | 51 | female | 1 | 1 |
| 46 | A | 4 | 3 | 32 | female | 0 | 0 |
| 47 | A | 11 | 3 | 45 | female | 0 | 0 |
| 48 | A | 11 | 3 | 64 | female | 0 | 1 |
| 49 | A | 11 | 3 | 46 | female | 1 | 0 |
| 50 | A | 11 | 3 | 65 | female | 0 | 0 |
| 51 | A | 11 | 3 | 54 | female | 0 | 0 |
| 52 | A | 11 | 3 | 55 | female | 0 | NA |
| 53 | B | 3 | 1 | 38 | female | 0 | 1 |
| 54 | B | 3 | 1 | 65 | female | 0 | 0 |
| 55 | B | 3 | 1 | 60 | female | 0 | 0 |
| 56 | B | 8 | 1 | 76 | female | 0 | 0 |
| 57 | B | 8 | 1 | 40 | female | 0 | 0 |
| 58 | B | 8 | 1 | 80 | female | 0 | 0 |
| 59 | B | 1 | 2 | 48 | female | 1 | 0 |
| 60 | B | 1 | 2 | 51 | female | 0 | 0 |
| 61 | B | 1 | 2 | 47 | female | 0 | 0 |
| 62 | B | 2 | 2 | 59 | female | 0 | 0 |
| 63 | B | 2 | 2 | 78 | female | 0 | 0 |
| 64 | B | 2 | 2 | 53 | female | 0 | 0 |
| 65 | B | 2 | 2 | 66 | female | 0 | 0 |
| 66 | B | 2 | 2 | 60 | female | 0 | 1 |
| 67 | B | 2 | 2 | 38 | female | 2 | NA |
| 68 | B | 2 | 2 | 51 | female | 0 | 0 |
| 69 | B | 2 | 2 | 73 | female | 0 | 1 |
| 70 | B | 4 | 2 | 48 | female | 0 | 1 |
| 71 | B | 4 | 2 | 51 | female | 0 | 0 |
| 72 | B | 4 | 2 | 34 | female | 0 | 1 |
| 73 | B | 4 | 2 | 67 | female | 0 | 0 |
| 74 | B | 4 | 2 | 55 | female | 0 | 1 |
| 75 | B | 4 | 2 | 57 | female | 0 | 1 |
| 76 | B | 4 | 2 | 53 | female | 1 | 1 |
| 77 | B | 5 | 2 | 41 | female | 0 | 0 |
| 78 | B | 5 | 2 | 42 | female | 0 | 0 |
| 79 | B | 5 | 2 | 39 | female | 2 | 0 |
| 80 | B | 5 | 2 | 38 | female | 0 | 0 |
| 81 | B | 7 | 2 | 59 | female | 0 | 2 |
| 82 | B | 7 | 2 | 35 | female | 0 | 0 |
| 83 | B | 7 | 2 | 34 | female | 0 | 0 |
| 84 | B | 7 | 2 | 56 | female | 0 | 0 |
| 85 | B | 7 | 2 | 45 | female | 0 | 1 |
| 86 | B | 7 | 2 | 53 | female | 0 | 0 |
| 87 | B | 7 | 2 | 52 | female | 0 | NA |
| 88 | B | 7 | 2 | 47 | female | 0 | 0 |
| 89 | B | 10 | 3 | 62 | female | 0 | 2 |
| 90 | B | 10 | 3 | 48 | female | 2 | 1 |
| 91 | B | 10 | 3 | 42 | female | 0 | 0 |
| 92 | B | 10 | 3 | 56 | female | 0 | 1 |
| 93 | B | 10 | 3 | 24 | female | 0 | 0 |
| 94 | B | 10 | 3 | 62 | female | 0 | 0 |
| 95 | B | 11 | 3 | 43 | female | 0 | 0 |
| 96 | B | 11 | 3 | 45 | female | 0 | 0 |
| 97 | B | 11 | 3 | 66 | female | 0 | 0 |
| 98 | B | 11 | 3 | 63 | female | 0 | 1 |
| 99 | B | 11 | 3 | 64 | female | 0 | 1 |
| 100 | B | 11 | 3 | 55 | female | 0 | 0 |
| 101 | B | 11 | 3 | 41 | female | 0 | 0 |
| 102 | B | 12 | 3 | 50 | female | 0 | 0 |
| 103 | B | 12 | 3 | 53 | female | 0 | 1 |
| 104 | B | 12 | 3 | 30 | female | 0 | 0 |
| 105 | B | 12 | 3 | 41 | female | 0 | 1 |
| 106 | B | 12 | 3 | 61 | female | 0 | 1 |

^a^BRCA1/2 status, 1 represents rare mutation and 2 represents pathogenic mutation. ^b^Numbers represent the count of lymph nodes, NA is short for not available.

**Supplementary Table 4 Age and sex of CTRL and their distribution in each type, class, and group.**

| **Index** | **Type** | **Class** | **Group** | **Age** | **Sex** |
| --- | --- | --- | --- | --- | --- |
| 1 | A | 1 | 3 | 80 | female |
| 2 | A | 6 | 2 | 74 | female |
| 3 | A | 13 | 2 | 76 | female |
| 4 | A | 12 | 2 | 74 | female |
| 5 | A | 10 | 2 | 78 | female |
| 6 | A | 1 | 3 | 70 | female |
| 7 | A | 1 | 3 | 71 | female |
| 8 | A | 14 | 1 | 71 | female |
| 9 | A | 6 | 2 | 70 | female |
| 10 | A | 12 | 2 | 70 | female |
| 11 | A | 5 | 2 | 82 | female |
| 12 | A | 7 | 1 | 73 | female |
| 13 | A | 11 | 3 | 81 | female |
| 14 | A | 15 | 2 | 80 | female |
| 15 | A | 8 | 1 | 72 | female |
| 16 | A | 5 | 2 | 76 | female |
| 17 | A | 9 | 2 | 74 | female |
| 18 | A | 15 | 2 | 75 | female |
| 19 | A | 7 | 1 | 82 | female |
| 20 | A | 10 | 2 | 79 | female |
| 21 | A | 15 | 2 | 71 | female |
| 22 | A | 5 | 2 | 75 | female |
| 23 | A | 1 | 3 | 72 | female |
| 24 | A | 13 | 2 | 81 | female |
| 25 | A | 15 | 2 | 78 | female |
| 26 | A | 5 | 2 | 83 | female |
| 27 | A | 6 | 2 | 70 | female |
| 28 | A | 10 | 2 | 77 | female |
| 29 | A | 13 | 2 | 70 | female |
| 30 | A | 8 | 1 | 74 | female |
| 31 | A | 3 | 3 | 81 | female |
| 32 | A | 2 | 3 | 72 | female |
| 33 | A | 10 | 2 | 72 | female |
| 34 | A | 4 | 3 | 75 | female |
| 35 | A | 15 | 2 | 96 | female |
| 36 | A | 10 | 2 | 88 | female |
| 37 | A | 1 | 3 | 87 | female |
| 38 | A | 9 | 2 | 81 | female |
| 39 | A | 14 | 1 | 80 | female |
| 40 | A | 9 | 2 | 80 | female |
| 41 | A | 8 | 1 | 86 | female |
| 42 | A | 8 | 1 | 76 | female |
| 43 | A | 13 | 2 | 88 | female |
| 44 | A | 13 | 2 | 90 | female |
| 45 | A | 12 | 2 | 88 | female |
| 46 | A | 11 | 3 | 88 | female |
| 47 | A | 5 | 2 | 82 | female |
| 48 | A | 5 | 2 | 83 | female |
| 49 | A | 13 | 2 | 87 | female |
| 50 | A | 2 | 3 | 79 | female |
| 51 | A | 7 | 1 | 76 | female |
| 52 | A | 2 | 3 | 85 | female |
| 53 | A | 7 | 1 | 83 | female |
| 54 | A | 2 | 3 | 84 | female |
| 55 | A | 6 | 2 | 77 | female |
| 56 | A | 7 | 1 | 77 | female |
| 57 | A | 1 | 3 | 79 | female |
| 58 | A | 13 | 2 | 75 | female |
| 59 | A | 12 | 2 | 80 | female |
| 60 | A | 10 | 2 | 80 | female |
| 61 | A | 13 | 2 | 73 | female |
| 62 | A | 6 | 2 | 80 | female |
| 63 | A | 15 | 2 | 83 | female |
| 64 | A | 10 | 2 | 87 | female |
| 65 | A | 13 | 2 | 78 | female |
| 66 | A | 12 | 2 | 83 | female |
| 67 | A | 6 | 2 | 86 | female |
| 68 | A | 10 | 2 | 81 | female |
| 69 | A | 9 | 2 | 80 | female |
| 70 | A | 13 | 2 | 85 | female |
| 71 | A | 13 | 2 | 87 | female |
| 72 | A | 7 | 1 | 85 | female |
| 73 | A | 15 | 2 | 83 | female |
| 74 | A | 6 | 2 | 93 | female |
| 75 | A | 9 | 2 | 81 | female |
| 76 | A | 5 | 2 | 77 | female |
| 77 | A | 12 | 2 | 79 | female |
| 78 | A | 7 | 1 | 72 | female |
| 79 | A | 6 | 2 | 84 | female |
| 80 | A | 9 | 2 | 84 | female |
| 81 | A | 7 | 1 | 77 | female |
| 82 | A | 5 | 2 | 94 | female |
| 83 | A | 5 | 2 | 76 | female |
| 84 | A | 6 | 2 | 83 | female |
| 85 | A | 13 | 2 | 62 | female |
| 86 | A | 14 | 1 | 83 | female |
| 87 | A | 12 | 2 | 81 | female |
| 88 | A | 13 | 2 | 81 | female |
| 89 | A | 13 | 2 | 85 | female |
| 90 | A | 10 | 2 | 95 | female |
| 91 | A | 8 | 1 | 88 | female |
| 92 | A | 13 | 2 | 81 | female |
| 93 | A | 10 | 2 | 81 | female |
| 94 | A | 10 | 2 | 73 | female |
| 95 | A | 10 | 2 | 86 | female |
| 96 | A | 10 | 2 | 88 | female |
| 97 | A | 14 | 1 | 85 | female |
| 98 | A | 13 | 2 | 84 | female |
| 99 | A | 6 | 2 | 84 | female |
| 100 | A | 13 | 2 | 75 | female |
| 101 | A | 10 | 2 | 80 | female |
| 102 | A | 13 | 2 | 91 | female |
| 103 | B | 9 | 1 | 70 | female |
| 104 | B | 8 | 1 | 75 | female |
| 105 | B | 8 | 1 | 74 | female |
| 106 | B | 11 | 3 | 79 | female |
| 107 | B | 10 | 3 | 71 | female |
| 108 | B | 8 | 1 | 83 | female |
| 109 | B | 1 | 2 | 75 | female |
| 110 | B | 2 | 2 | 73 | female |
| 111 | B | 8 | 1 | 73 | female |
| 112 | B | 4 | 2 | 75 | female |
| 113 | B | 4 | 2 | 75 | female |
| 114 | B | 11 | 3 | 73 | female |
| 115 | B | 9 | 1 | 75 | female |
| 116 | B | 13 | 1 | 77 | female |
| 117 | B | 13 | 1 | 71 | female |
| 118 | B | 10 | 3 | 79 | female |
| 119 | B | 9 | 1 | 84 | female |
| 120 | B | 8 | 1 | 76 | female |
| 121 | B | 9 | 1 | 70 | female |
| 122 | B | 9 | 1 | 85 | female |
| 123 | B | 10 | 3 | 71 | female |
| 124 | B | 8 | 1 | 71 | female |
| 125 | B | 7 | 2 | 72 | female |
| 126 | B | 8 | 1 | 73 | female |
| 127 | B | 10 | 3 | 74 | female |
| 128 | B | 7 | 2 | 79 | female |
| 129 | B | 8 | 1 | 72 | female |
| 130 | B | 2 | 2 | 79 | female |
| 131 | B | 8 | 1 | 78 | female |
| 132 | B | 8 | 1 | 73 | female |
| 133 | B | 1 | 2 | 70 | female |
| 134 | B | 11 | 3 | 70 | female |
| 135 | B | 2 | 2 | 73 | female |
| 136 | B | 13 | 1 | 73 | female |
| 137 | B | 9 | 1 | 71 | female |
| 138 | B | 7 | 2 | 74 | female |
| 139 | B | 8 | 1 | 70 | female |
| 140 | B | 7 | 2 | 75 | female |
| 141 | B | 3 | 1 | 97 | female |
| 142 | B | 11 | 3 | 88 | female |
| 143 | B | 12 | 3 | 86 | female |
| 144 | B | 11 | 3 | 80 | female |
| 145 | B | 1 | 2 | 88 | female |
| 146 | B | 9 | 1 | 90 | female |
| 147 | B | 8 | 1 | 78 | female |
| 148 | B | 10 | 3 | 86 | female |
| 149 | B | 11 | 3 | 72 | female |
| 150 | B | 6 | 2 | 89 | female |
| 151 | B | 10 | 3 | 84 | female |
| 152 | B | 9 | 1 | 92 | female |
| 153 | B | 3 | 1 | 82 | female |
| 154 | B | 6 | 2 | 79 | female |
| 155 | B | 10 | 3 | 86 | female |
| 156 | B | 9 | 1 | 97 | female |
| 157 | B | 9 | 1 | 75 | female |
| 158 | B | 13 | 1 | 81 | female |
| 159 | B | 4 | 2 | 83 | female |
| 160 | B | 13 | 1 | 91 | female |
| 161 | B | 10 | 3 | 80 | female |
| 162 | B | 2 | 2 | 81 | female |
| 163 | B | 8 | 1 | 84 | female |
| 164 | B | 7 | 2 | 74 | female |
| 165 | B | 7 | 2 | 84 | female |
| 166 | B | 6 | 2 | 81 | female |
| 167 | B | 10 | 3 | 82 | female |
| 168 | B | 5 | 2 | 79 | female |
| 169 | B | 13 | 1 | 86 | female |
| 170 | B | 7 | 2 | 79 | female |
| 171 | B | 9 | 1 | 86 | female |
| 172 | B | 2 | 2 | 66 | female |
| 173 | B | 13 | 1 | 91 | female |
| 174 | B | 3 | 1 | 91 | female |
| 175 | B | 9 | 1 | 84 | female |
| 176 | B | 4 | 2 | 80 | female |
| 177 | B | 2 | 2 | 79 | female |
| 178 | B | 8 | 1 | 77 | female |
| 179 | B | 8 | 1 | 85 | female |
| 180 | B | 6 | 2 | 71 | female |
| 181 | B | 8 | 1 | 82 | female |
| 182 | B | 7 | 2 | 88 | female |
| 183 | B | 7 | 2 | 79 | female |
| 184 | B | 9 | 1 | 73 | female |
| 185 | B | 7 | 2 | 78 | female |
| 186 | B | 10 | 3 | 83 | female |
| 187 | B | 11 | 3 | 88 | female |
| 188 | B | 13 | 1 | 81 | female |
| 189 | B | 13 | 1 | 83 | female |
| 190 | B | 6 | 2 | 94 | female |
| 191 | B | 8 | 1 | 88 | female |
| 192 | B | 11 | 3 | 90 | female |
| 193 | B | 10 | 3 | 91 | female |
| 194 | B | 13 | 1 | 89 | female |
| 195 | B | 7 | 2 | 86 | female |
| 196 | B | 11 | 3 | 91 | female |
| 197 | B | 2 | 2 | 92 | female |
| 198 | B | 1 | 2 | 93 | female |
| 199 | B | 7 | 2 | 85 | female |
| 200 | B | 10 | 3 | 73 | female |
| 201 | B | 4 | 2 | 87 | female |
| 202 | B | 9 | 1 | 72 | female |
| 203 | B | 6 | 2 | 66 | female |
| 204 | B | 9 | 1 | 78 | female |
| 205 | B | 12 | 3 | 82 | female |
| 206 | B | 10 | 3 | 81 | female |
| 207 | B | 8 | 1 | 80 | female |
| 208 | B | 4 | 2 | 84 | female |
| 209 | B | 3 | 1 | 94 | female |
| 210 | B | 10 | 3 | 83 | female |
| 211 | B | 1 | 2 | 81 | female |
| 212 | B | 7 | 2 | 78 | female |
| 213 | B | 4 | 2 | 79 | female |
| 214 | B | 10 | 3 | 84 | female |
| 215 | B | 13 | 1 | 46 | female |
| 216 | B | 8 | 1 | 90 | female |
| 217 | B | 4 | 2 | 67 | female |
| 218 | B | 7 | 2 | 77 | female |
| 219 | B | 8 | 1 | 73 | female |
| 220 | B | 8 | 1 | 92 | female |
| 221 | B | 8 | 1 | 83 | female |
| 222 | B | 3 | 1 | 76 | female |
| 223 | B | 7 | 2 | 79 | female |
| 224 | B | 2 | 2 | 81 | female |
| 225 | B | 3 | 1 | 88 | female |
| 226 | B | 4 | 2 | 85 | female |
| 227 | B | 13 | 1 | 72 | female |
| 228 | B | 13 | 1 | 70 | female |
| 229 | B | 9 | 1 | 79 | female |
| 230 | B | 4 | 2 | 75 | female |
| 231 | B | 6 | 2 | 82 | female |
| 232 | B | 12 | 3 | 66 | female |
| 233 | B | 13 | 1 | 75 | female |
| 234 | B | 10 | 3 | 77 | female |
| 235 | B | 6 | 2 | 78 | female |
| 236 | B | 9 | 1 | 78 | female |
| 237 | B | 5 | 2 | 81 | female |
| 238 | B | 5 | 2 | 81 | female |
| 239 | B | 1 | 2 | 84 | female |
| 240 | B | 8 | 1 | 86 | female |
| 241 | B | 7 | 2 | 68 | female |
| 242 | B | 1 | 2 | 82 | female |
| 243 | B | 13 | 1 | 75 | female |
| 244 | B | 8 | 1 | 71 | female |
| 245 | B | 10 | 3 | 81 | female |
| 246 | B | 5 | 2 | 82 | female |
| 247 | B | 4 | 2 | 76 | female |
| 248 | B | 3 | 1 | 80 | female |
| 249 | B | 1 | 2 | 86 | female |
| 250 | B | 2 | 2 | 71 | female |
| 251 | B | 4 | 2 | 85 | female |
| 252 | B | 5 | 2 | 88 | female |
| 253 | B | 8 | 1 | 80 | female |
| 254 | B | 7 | 2 | 81 | female |
| 255 | B | 5 | 2 | 89 | female |
| 256 | B | 8 | 1 | 77 | female |
| 257 | B | 7 | 2 | 88 | female |
| 258 | B | 7 | 2 | 85 | female |
| 259 | B | 12 | 3 | 86 | female |
| 260 | B | 5 | 2 | 84 | female |
| 261 | B | 9 | 1 | 78 | female |
| 262 | B | 5 | 2 | 83 | female |
| 263 | B | 7 | 2 | 75 | female |
| 264 | B | 10 | 3 | 84 | female |
| 265 | B | 5 | 2 | 80 | female |
| 266 | B | 10 | 3 | 70 | female |
| 267 | B | 9 | 1 | 97 | female |
| 268 | B | 5 | 2 | 87 | female |
| 269 | B | 7 | 2 | 80 | female |
| 270 | B | 9 | 1 | 81 | female |
| 271 | B | 7 | 2 | 87 | female |
| 272 | B | 13 | 1 | 83 | female |
| 273 | B | 6 | 2 | 72 | female |
| 274 | B | 3 | 1 | 87 | female |
| 275 | B | 9 | 1 | 74 | female |
| 276 | B | 7 | 2 | 75 | female |
| 277 | B | 11 | 3 | 96 | female |
| 278 | B | 9 | 1 | 71 | female |
| 279 | B | 9 | 1 | 86 | female |
| 280 | B | 13 | 1 | 85 | female |
| 281 | B | 11 | 3 | 78 | female |
| 282 | B | 6 | 2 | 91 | female |
| 283 | B | 13 | 1 | 84 | female |
| 284 | B | 7 | 2 | 84 | female |
| 285 | B | 10 | 3 | 88 | female |
| 286 | B | 7 | 2 | 97 | female |
| 287 | B | 9 | 1 | 71 | female |
